# Structural dynamics at cytosolic inter-protomer interfaces control gating of a mammalian TRPM5 channel

**DOI:** 10.1101/2023.08.26.554973

**Authors:** Sebastian Karuppan, Lynn Goss Schrag, Andrés Jara-Oseguera, Lejla Zubcevic

**Affiliations:** Department of Biochemistry and Molecular Biology, The University of Kansas Medical Center, Kansas City, KS, 66160, USA; Department of Molecular Biosciences, College of Natural Sciences, The University of Texas at Austin, TX, 78712, USA

**Author notes:** Correspondence to: Andrés Jara-Oseguera,; Lejla Zubcevic. These authors contributed equally (Sebastian Karuppan, Lynn Goss Schrag). Biodesign Institute, Arizona State University, Tempe, AZ, 85281, USA.

## Abstract

The Transient Receptor Potential Melastatin (TRPM) family of tetrameric cation channels is involved in a wide range of biological functions, from sensory perception to cardiac function. The structurally conserved TRPM cytoplasmic domains make up > 70 % of the total protein. To investigate the mechanism by which the TRPM cytoplasmic domains contribute to gating, we employed electrophysiology and cryo-EM to study TRPM5 – a channel that primarily relies on activation via intracellular Ca^2+^. Here we show that activation of mammalian TRPM5 channels is strongly altered by desensitization. Structures of rat TRPM5 identify a series of conformational transitions triggered by Ca^2+^ binding. These transitions involve formation and dissolution of cytoplasmic inter-protomer interfaces, where the MHR1/2 domain and the rib helix play important roles. This study shows the importance of the cytoplasmic assembly in TRPM5 channel function and sets the stage for future investigations of other members of the TRPM family.

## Introduction

The Transient Receptor Potential Melastatin (TRPM) family of tetrameric cation channels encompasses eight members (TRPM1-TRPM8) that are involved in a wide range of biological functions, from temperature sensing and taste transduction^1–4^ to regulation of cardiac function^5^, inflammatory pain and insulin secretion^6–8^. In recent years, cryo-EM has contributed greatly to our understanding of these ion channels, and we now have representative structures from almost every TRPM family member^9–11^. One of their distinguishing features is a very large cytoplasmic domain, which makes up > 70 % of the total protein and is fairly structurally conserved throughout the subfamily. It consists of three amino (N) - terminal Melastatin Homology Repeat domains (MHR1/2, MHR3, and MHR4) that are arranged around a C-terminal umbrella-like structure made up of four rib helices that run nearly parallel with the membrane plane, extending from a central coiled coil domain with an axis perpendicular to the membrane. Mutations in the cytoplasmic assembly of TRPMs can lead to severe disease, including cardiac conductance block^12–14^ and complete congenital stationary night blindness^15,16^, suggesting a critical role of these domains in channel function.

To determine the mechanism by which the TRPM cytoplasmic domains contribute to gating in TRPM channels, we decided to focus on TRPM5 – a monovalent cation-selective channel^17^ that primarily relies on activation by cytoplasmic Ca^2+^ ions^17,18^. The TRPM5 channel is important for signaling in type II taste bud cells^19,20^, and for insulin secretion in pancreatic β-cells^21 6,7,21–24^. Because of this, TRPM5 is viewed as an attractive pharmacological target for the treatment of type II diabetes^25^. Structures of zebrafish TRPM5 were published recently^11^, and the study identified two Ca^2+^-binding sites within the channel: one in the Voltage Sensing-Like Domain (VSLD) within the transmembrane domain and the other in the cytoplasmic domain, at the interface between MHR1/2 and MHR3. The binding site in the VSLD is conserved amongst many TRPM channels, including TRPM4 and TRPM8^26–30^, but the intracellular binding site is unique to TRPM5. Both Ca^2+^-binding sites in TRPM5 are highly conserved across species^11^. However, there is otherwise limited sequence identity between the zebrafish TRPM5 channel and its mammalian orthologues (∼ 57% sequence identity between the zebrafish channel and the rat or human TRPM5 orthologs). We therefore set out to determine the molecular underpinnings of mammalian TRPM5 activation by Ca^2+^.

Using a combined approach of cryo-EM and electrophysiology, here we show that gating of the rat, human, and mouse TRPM5 orthologues is functionally distinct from zebrafish and that the cytoplasmic domains play an important role in mediating Ca^2+^-dependent activation and desensitization. We observe novel structural dynamics associated with mammalian TRPM5 cytoplasmic domains and interfaces and provide evidence for their contribution to channel function that might also be relevant for other TRPM channels.

## Results

### Mammalian TRPM5 channels have a distinct structural and functional profile

We began by assessing current responses to intracellular Ca^2+^ in inside-out patches excised from HEK293 cells expressing human, rat, mouse, or zebrafish TRPM5 channels as well as control cells expressing only GFP (Fig. 1). Although we detected a small, desensitizing endogenous current activated by μM Ca^2+^ concentrations in GFP-only controls (Fig. 1a), TRPM5-mediated currents were clearly distinguishable from background by their robust outward-rectifying responses to nanomolar Ca^2+^ that were completely absent in the GFP-only controls (Fig. 1b-g). These currents were similar for all four TRPM5 orthologues tested. However, responses to micromolar Ca^2+^ in the three mammalian channels (Fig. 1b-d) were drastically different from zebrafish (Fig. 1e): upon exposure to 5.95 micromolar Ca^2+^, currents reached a peak of similar amplitude at positive and negative membrane voltages, followed by a rapid decay in current that was much more pronounced in the human, rat, and mouse channels than in the zebrafish orthologue (Fig. 1f). This marked difference in the extent of desensitization underscores a key difference in gating between mammalian and zebrafish TRPM5 channels.

**Fig. 1.**
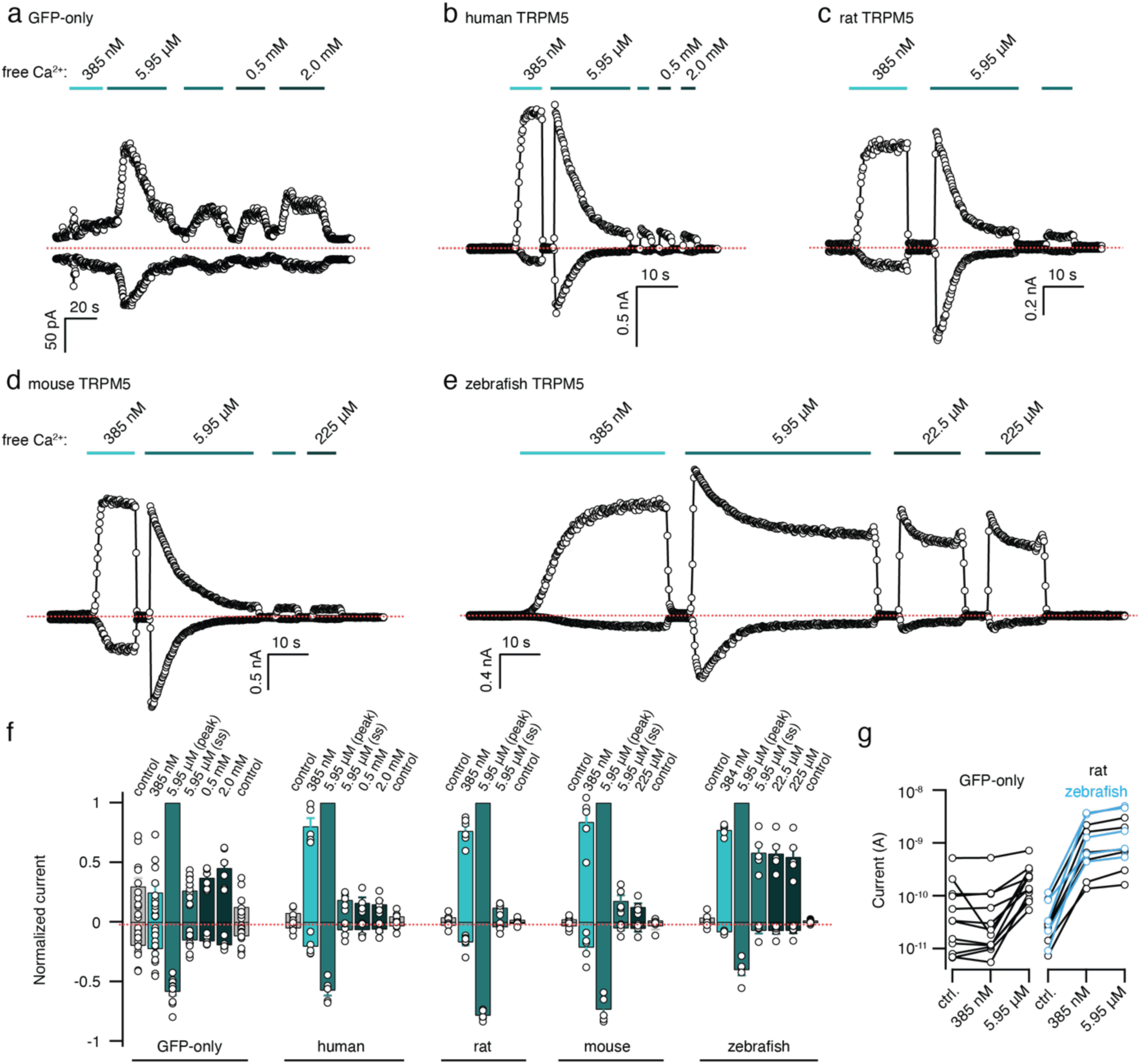
Mammalian and zebrafish TRPM5 channels possess a distinct activation profile. **a-e** Representative current time-courses elicited by pulses from 0 to ± 140 mV on inside-out patches obtained from transiently transfected HEK293 cells exposed to intracellular solutions containing distinct free Ca^2+^ concentrations denoted by the horizontal bars. Dotted lines denote the zero-current level. **f** Mean ± SEM currents obtained from experiments as in (a-e), normalized to the peak current at +140 mV in the presence of 5.95 μM free Ca^2+^. Data from individual experiments are shown as circles (n = 5-12). **g** Current magnitudes at +140 mV from individual patches obtained from cells transfected with GFP only, or GFP together with rat (black) or zebrafish (blue) TRPM5 channel cDNA (as in a, c and e). For each patch, data is shown in control solution, and in the presence of 385 nM (steady-state) or 5.95 μM (peak) free Ca^2+^.

In desensitized channels activation by Ca^2+^ was retained but diminished in amplitude even in the presence of much larger concentrations of the cation (Fig. 1f), as previously observed^18^. We carefully measured the concentration of free Ca^2+^ in our recording solutions using Fura-2 (Extended Data Fig. 1a), and obtained dose-response relations to Ca^2+^ for the rat TRPM5 channel before and after complete desensitization (Extended Data Fig. 1b-e). Activation by Ca^2+^ was extremely cooperative, suggesting the involvement of at least two Ca^2+^ sites per subunit. In addition, desensitization did not largely alter the EC_50_ for Ca^2+^ activation nor the apparent cooperativity, but strongly reduced current amplitudes, particularly at negative voltages (Extended Data Fig. 1b-e). We also found that the rate of desensitization depended steeply on Ca^2+^ at concentrations close to the EC_50_ for channel activation (Extended Data Fig. 1f-h), indicating that activation and desensitization are tightly coupled to cooperative Ca^2+^ binding to multiple sites per channel subunit. Finally, we found that very large quantities (200 μM) of diC8 PIP_2_, a soluble analogue of the positive channel modulator phosphoinositide-4,5-bisphosphate, were sufficient to almost completely restore desensitized currents to their pre-desensitization values at high Ca^2+^ concentrations (Extended Data Fig. 2).

Together, our results indicate that mammalian TRPM5 channels undergo multiple conformational transitions following Ca^2+^-binding, involving energetically distinct activation pathways before and after desensitization, and that desensitization likely arises from Ca^2+^- dependent conformational changes in the channel as well as PIP_2_ depletion from the membrane. We therefore set out to determine the molecular mechanisms that underlie Ca^2+^-dependent activation and desensitization of rTRPM5. First, to investigate the conformation that the rTRPM5 channel occupies in the absence of stimulus, we expressed and purified the rTRPM5 channel and determined its cryo-EM structure in the absence of Ca^2+^ (rTRPM5_EGTA_, Extended Data Fig. 3). Consistently, an inspection of the previously reported locations for Ca^2+^ binding^11^ indicated that no Ca^2+^ ions were present in any of the eight binding sites within the rTRPM5_EGTA_ tetramer (Extended Data Fig. 4a-h).

The rTRPM5_EGTA_ structure possesses all the canonical elements of a TRPM channel (Extended Data Fig. 5a). Namely, it assembles as a tetramer and each protomer is composed of an N-terminal domain consisting of MHR1/2, MHR3, and MHR4, and a coupling domain (CD) between the cytoplasmic and transmembrane domains (TM). The TM domain is made up of a VSLD bundle (S1-S4 helices) and a pore domain (S5-S6 helices) which are linked by a short helical S4-S5 linker. The C-terminal TRP domain is nestled by the VSLD on top and the helix-loop-helix (HLH) motif of the coupling domain (CD) on the bottom. From the TRP domain, a short helix extends down into the cytoplasm and links up to the rib helix which runs parallel with the membrane plane. The rib helix then connects to a coiled coil structure at a ∼120° angle. (Extended Data Fig. 5a). The MHR1/2 domain is made up of nine helices sandwiching a β sheet (Extended Data Fig. 5b) and the MHR3 domain is formed by a stack of helices arranged in a helix-turn-helix (HTH) manner to produce a spring-like structure (Extended Data Fig. 5c).

The rTRPM5_EGTA_ tetrameric channel adopts a unique conformation as it assembles with near-C4-fold symmetry in the TM domain but shows no symmetry (C1) in the cytoplasmic domain.

The C1 symmetry is most pronounced in the parts of the channel located deepest in the cytoplasm: the MHR1/2 and the distal parts of the coiled coil (Fig. 2a, Extended Data Fig. 3). This leads to an arrangement at the cytoplasmic side of the channel where each interprotomer interface is unique (Fig. 2b-c). Namely, the distance between the MHR domain and the rib helix of neighboring protomers is different at each interface – ranging from 22 Å to 15 Å when measured from Cα of K94 in the MHR1/2 to Cα of N1090 in the rib helix (Fig. 2c).

**Fig. 2.**
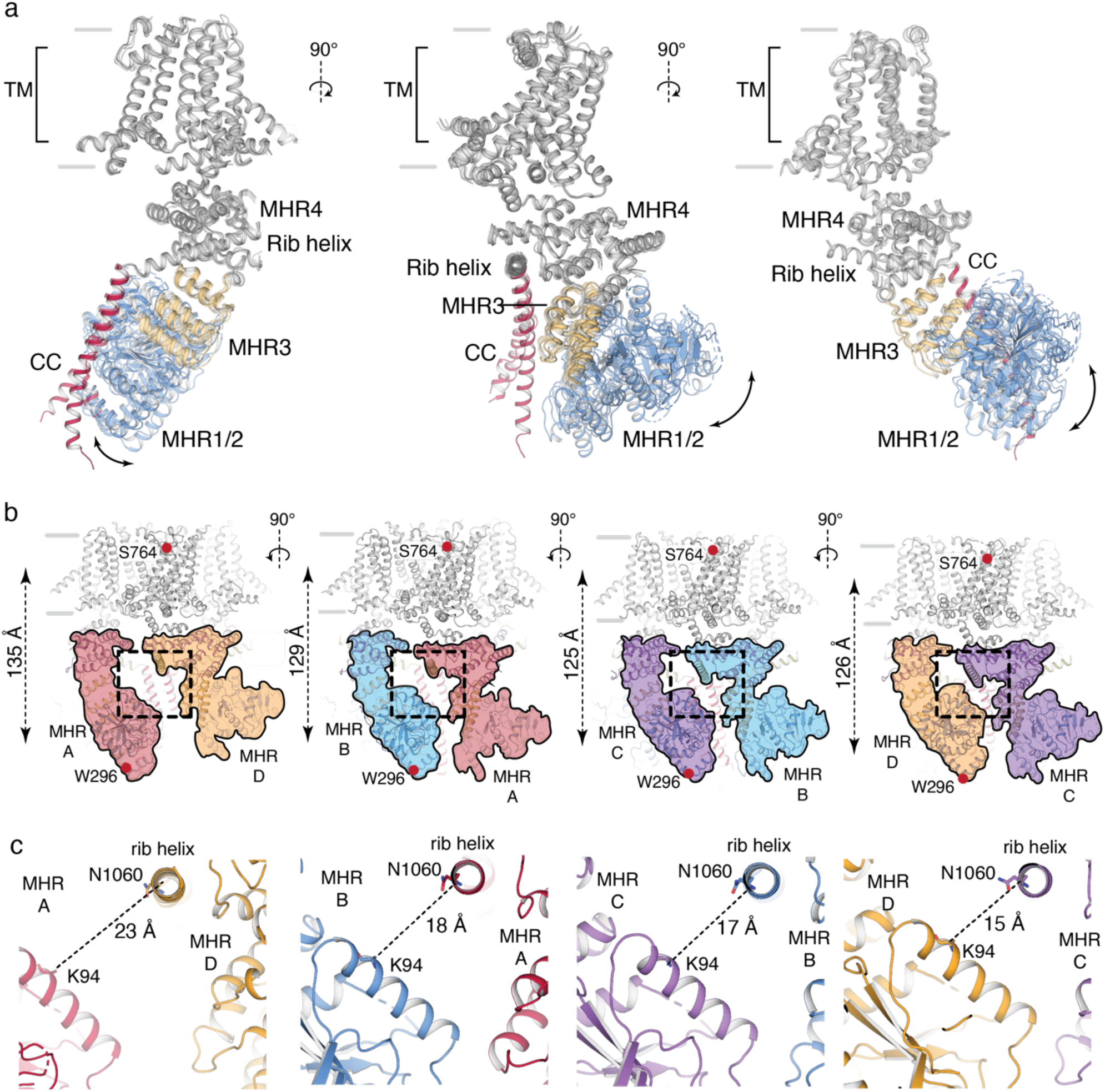
Structure of the rTRPM5 in the presence of EGTA. **a** Structure of the rat TRPM5 channel in the presence of EGTA is asymmetric. Individual protomers are overlaid to show that the asymmetry is most pronounced in the cytoplasmic domains, and especially in the MHR1/2 domain (blue cartoon) and the coiled coil (CC, red cartoon). **b** Each cytoplasmic inter-protomer interface is unique in rTRPM5_EGTA_. The MHR domains of each protomer are shaded differently (A: red; B: marine; C: purple; D: gold). The MHR domains of protomer A (red) extend deep into the cytosol, so that the dimensions of the channel from the top of the transmembrane domain (S764) to the bottom of the MHR domain (W296) are ∼ 135 Å. The MHR domains in protomers B (purple) and D (gold) appear to rotate upwards, which shortens the length of the channel to ∼ 125 Å as well as the distance to the neighboring protomers. **c** Close-up of the boxed region in **b**. The conformation of the MHR domains also dictates the distance between protomers. In protomer A (red), where the MHR domains are extended into the cytosol, the distance to the neighboring protomer D is large (23 Å between the Cα of K94 of MHR A and N1090 of MHR D). This distance shortens upon rotation of the MHR domains, as observed in MHR B, C, and D.

It is worth noting that we observe a non-protein density in the rTRPM5_EGTA_ map which may correspond to PIP_2_. This density occupies a cleft created by the pre-S1 domain, helices S1 and S4, the S4-S5 linker, the TRP domain, and the CD and overlaps significantly with the PIP_2_ binding site described in mouse TRPM8 (Extended Data Fig. 6)^30,31^. Consistent with the observation that the TM domain maintains C4 symmetry, this density is observed in all four protomers (Extended Data Fig. 6a-d). As no PIP_2_ was added to our sample prior to freezing, we assume that the lipid co-purified with the channel. Interestingly, PIP_2_ – a positive allosteric regulator for both TRPM8^32^ and TRPM5^18^ channels – was also found to co-purify with the mouse TRPM8^31^.

### The distal coiled coil and MHR domains are highly flexible in the absence of Ca^2+^

To determine the source of the asymmetry, we examined the cytoplasmic regions of rTRPM5_EGTA_. The bottom view of the channel showed that an interface exists between the coiled coil (CC) helix and the MHR1/2 domain of the neighboring protomer and that each one of these four CC-MHR1/2 interfaces is different (Fig. 3a) in the tetramer. A closer look at the CC structure in rTRPM5_EGTA_ offered an explanation to these observations. The CC contains a double glycine motif (GG hinge) that introduces a flexible point into this rigid assembly of helices (Fig. 3b) and divides the CC into two parts – the channel-proximal part, located above the GG hinge, and the distal part, located below (Fig. 3b). The distal part appears to be highly flexible and can assume various conformations. An overlay of the CC helices indicates that they begin to diverge below the GG hinge (Fig. 3c). Two of the CC helices from protomers B and C (CC B and CC C) are resolved for up to 7 helical turns beyond the GG. However, their conformations are different – one is straight and the other begins to curve at the GG hinge. The other two helices reflect higher flexibility: CC D is unresolved beyond the GG (presumably due to high flexibility) and CC A appears to be captured in the process of unwinding (Fig. 3b-c).

**Fig. 3.**
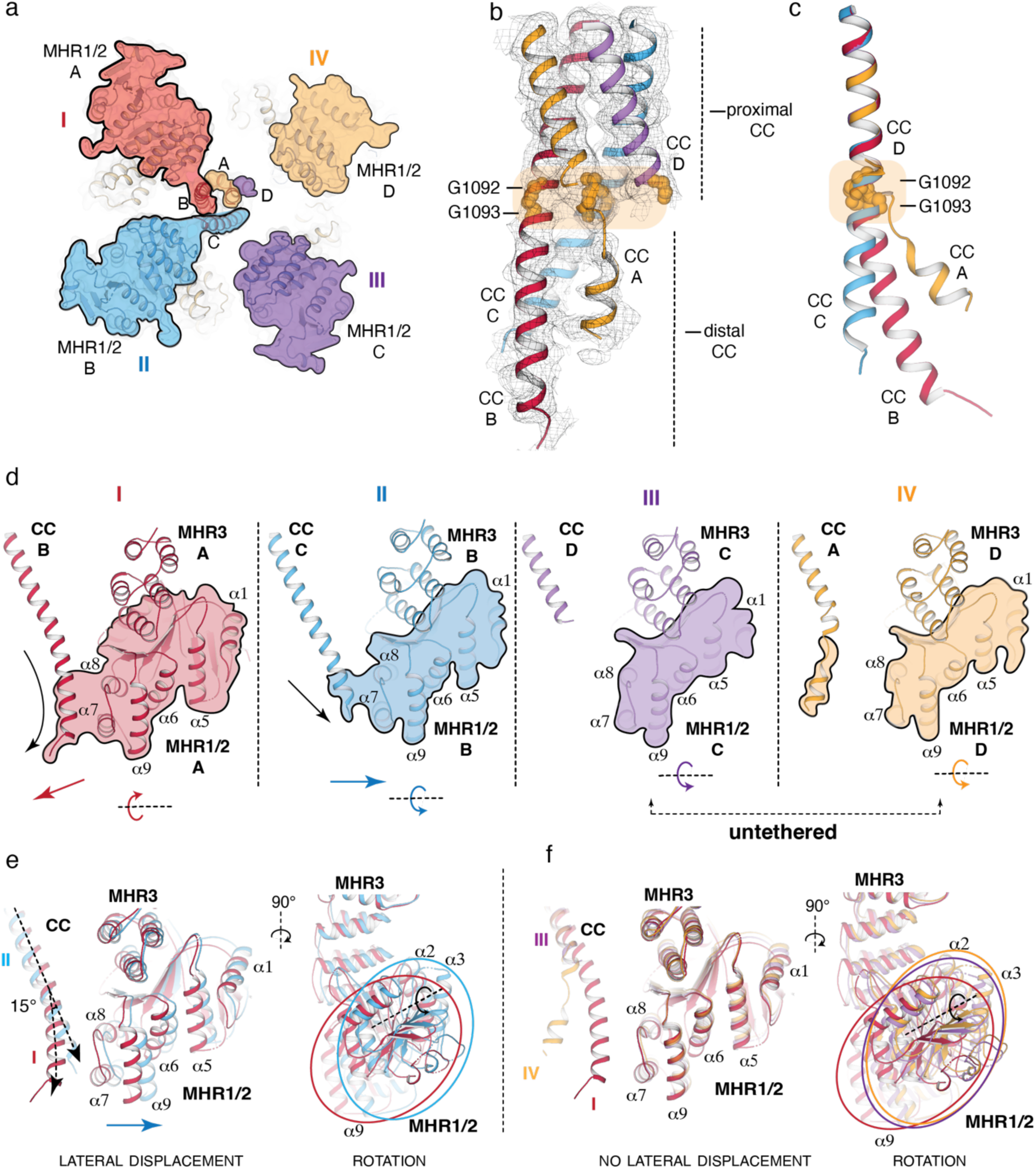
The distal CC controls the conformation of the MHR1/2 domain from its neighboring protomer. **a** Bottom-up view of the rat TRPM5 channel determined in EGTA. Four distinct interfaces are observed between the MHR1/2 domain and the CC of neighboring protomers in this structure. They are colored in red (interface I, MHR A and CC B), blue (interface II, MHR B and CC C), purple (interface III, MHR C and CC D), and gold (interface IV, MHR D and CC A). **b** The distinct interfaces are caused by the arrangement of the CC. The proximal part adopts a tetrameric coiled coil conformation, but the distal part (beyond the flexible GG hinge, shown in orange spheres and orange box) is highly asymmetric. The density for the CC assembly is rendered as mesh and shown at contour level 0.15. **c** An overlay of the individual CC helices shows that the asymmetry begins at the flexible GG hinge. **d** The distal CC dictates the conformation of the MHR1/2 domain from the neighboring protomer. Side view of interfaces I-IV are shown. For clarity, only the CC and MHR1/2 domains are shown. The long, curved helix of CC B pulls the MHR1/2 down and forces a forward rotation. The straight CC C pushes the MHR1/2 B laterally and forces a backward rotation. CC D and CC A form no interactions with their respective MHR1/2 domains due to being unresolved (CC D) or broken (CC A) beyond the GG hinge. The untethered MHR1/2 domains rotate backwards. **e** Overlay of interfaces I (red) and II (blue). In interface II, the CC pushes laterally on the MHR1/2 (lateral displacement visible in the front view (left panel)) and a forces a backward (counterclockwise) rotation around its internal β-sheet axis (right panel). The outline of the MHR1/2 of interface I is indicated with a red oval and the outline of MHR1/2 of interface II is indicated with a blue oval. The MHR1/2 helices α1- α3 and α5- α9 are labeled. The dashed line indicates the β-sheet axis and the arrow indicates the rotation. **f** Overlay of interfaces I (red), III (purple), and IV (gold). The MHR1/2 domains of III and IV are not subject to a lateral push from the CC (left panel), but untethered they rotate backward (counterclockwise) around their internal β-sheet axis (right panel). The outline of the MHR1/2 of interface I is indicated with a red oval, MHR1/2 of interface III with a purple oval, and MHR1/2 of interface IV with a gold oval. The MHR1/2 helices α1- α3 and α5- α9 are labeled. The dashed line indicates the β-sheet axis and the arrow indicates the rotation.

We next set out to determine how the interactions with the distal CC influence the conformation of the cytoplasmic interprotomer interfaces, and specifically, the MHR1/2 domain of the neighboring protomer. As mentioned earlier, the bottom-up view of the channel shows that each CC-MHR1/2 interface is unique. For ease of explanation, we termed these interfaces I to IV (Fig. 3a and d). Interface I consists of the CC of protomer B (CC B) and the MHR1/2 domains of protomer A (Fig. 3d, shown in red). This interface has the most extensive CC-MHR1/2 interactions because CC B is the longest of the four CC helices and because it possesses a curve which enables it to reach around and contact helices α7 and α8 of the neighboring MHR1/2 A (Fig. 3c-d, Extended Data Fig. 3h). The interaction with CC B pulls the MHR1/2 A deep into the cytosol. This elongates the channel itself, so that the distance between the top of S2 (V764) and bottom of MHR1/2 (W296) measures 135 Å, the longest of the four protomers (Fig. 2b).

The interface II is formed by CC C and MHR1/2 B. Here, the CC C helix adopts a straighter and seemingly more rigid conformation than CC B (Fig. 3c, shown in blue). Because CC C does not possess a curve, it appears to exert a lateral push on the MHR1/2 B (Figs. 3d-e, Extended Data Fig. 3i). This push from CC C also appears to rotate the MHR1/2 B domain around its internal β sheet axis – a movement that brings the MHR1/2 B domain closer to the neighboring protomer A (Figs. 3e, 2b) and compacts the length of the channel so that it measures 129 Å from the top of S2 (V764) and bottom of MHR1/2 (W296) (Fig. 2b).

At interfaces III and IV we do not observe any contacts between CC and MHR1/2 because these CC helices are either not resolved beyond the GG hinge (CC D) or are apparently unwinding (CC A) (Fig. 3b-d, Extended Data Fig. 3j-k). In both cases this leaves the MHR1/2 domains untethered from the central CC assembly. The untethered MHR1/2 domains rotate around their internal β sheet axis and swing towards the neighboring protomer (Figs. 2c and 3f). This rotation shortens the channel further (125-126 Å) from the top of S2 to the bottom of MHR1/2 (Fig. 2b).

Together these data indicate that the interface between the distal CC and the MHR1/2 domain of the neighboring subunit is highly dynamic, with two observed configurations: (1) An ordered, extended distal CC that interacts with the neighboring MHR1/2 and prevents it from interacting with the neighboring protomer and (2) a disordered (or highly flexible) distal CC that no longer contacts the MHR1/2 that thus swings sideways towards the neighboring subunit.

Conversely, these neighboring MHR domain interfaces in the zebrafish TRPM5 channel obtained in the presence of EDTA (PDB ID 7MBP)^11^ are tightly coupled (Extended Data Fig. 7a), as evidenced by a tight network of interactions between the MHR1/2 and the rib helix and MHR3 of the neighboring protomer (Extended Data Fig. 7e). Furthermore, the channel has C4 symmetry. The CC of the zebrafish channel is resolved only two helical turns beyond its GG hinge (Extended Data Fig. 7c) and therefore we observe no interactions between the CC and the MHR1/2 of the neighboring protomer. This is in stark contrast to the rTRPM5_EGTA_, which has 4 distinct – apparently uncoupled – cytoplasmic interprotomer interfaces (Fig. 2c). However, the differences between the cytoplasmic domains of rTRPM5_EGTA_ and the zebrafish TRPM5 in EDTA appear to have no bearing for the conformation of the pore, as the two channels align well in the TM regions (Extended Data Fig. 7b). In addition, the cytoplasmic assemblies in zebrafish structures in the presence of Ca^2+^ all appear to be strongly coupled (Extended Data Fig. 7f). This suggests that the mammalian channel has a more flexible cytoplasmic assembly than its zebrafish counterpart. It also suggests that non-concerted conformational dynamics at each of the four cytoplasmic domains observed in the absence of Ca^2+^ in the rTRPM5_EGTA_ structure are not strongly coupled to the conformation of the transmembrane domain. To test this, we introduced single and double alanine substitutions into the GG hinge (Fig. 4), and obtained responses to Ca^2+^ that were indistinguishable from WT. These three mutations are predicted to reduce flexibility at the CC. To test for the opposite effect, we deleted the distal CC portion of the C-terminal and obtained a similar result: channels with the deletion responded to Ca^2+^ similarly to WT (Fig. 4b-c). Together, these results indicate that the cytoplasmic assembly in mammalian TRPM5 channels is highly flexible in the absence of Ca^2+^, and that the distal coiled coil does not contribute to the energetics of Ca^2+^- dependent gating. To determine the conformational changes that occur upon agonist binding, we determined the structure of rTRPM5 in the presence of Ca^2+^.

**Fig. 4.**
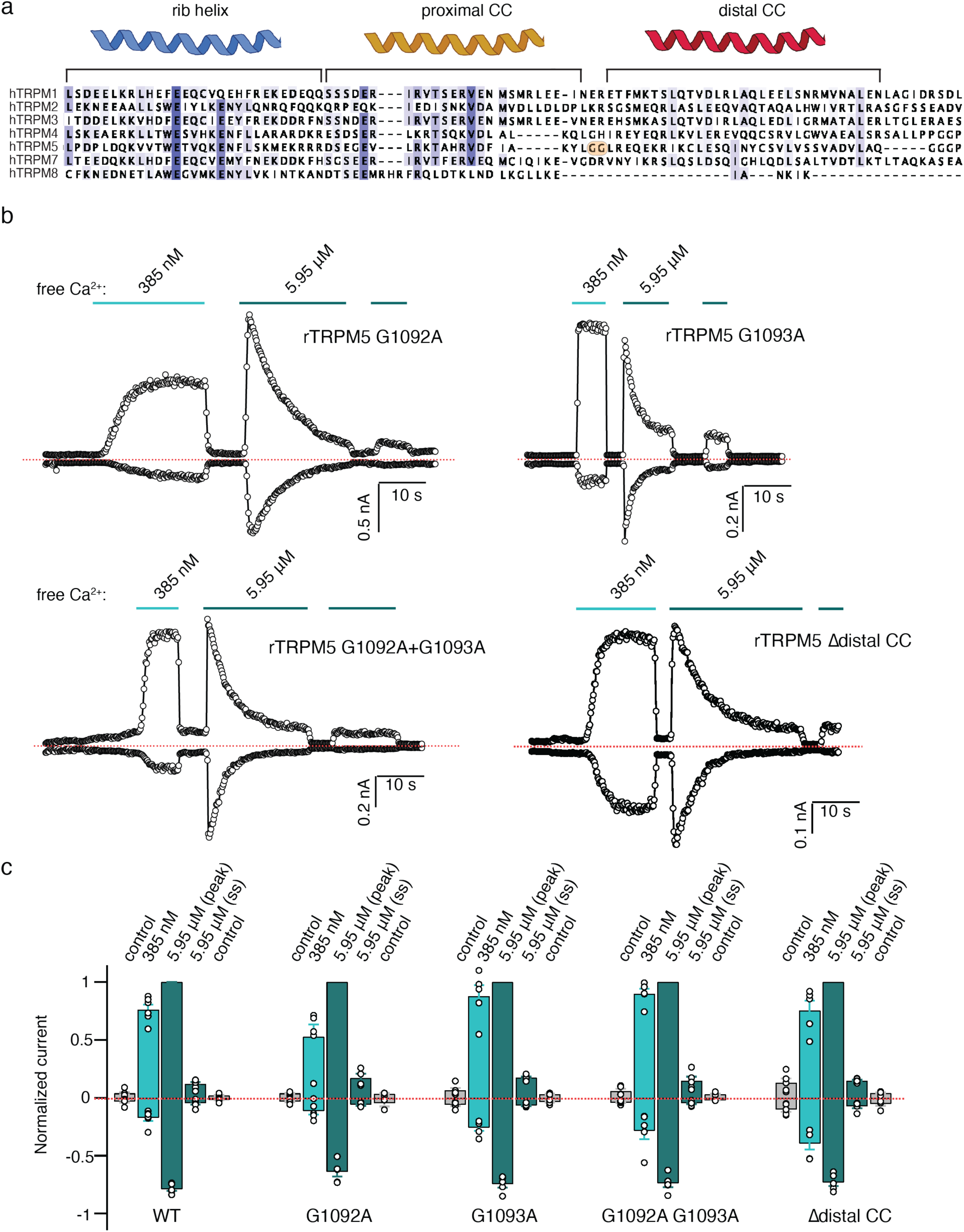
Glycine hinge mutants do not significantly affect activation and desensitization of rTRPM5 by Ca^2+^. **a** A sequence alignment of the rib helix, and the proximal and distal CC of human TRPM channels. Sequences of rat and zebrafish TRPM5 are included too. The GG hinge is marked with an orange box. **b** Representative time-courses at ± 140 mV obtained from inside-out patches from cells expressing rTRPM5 channel mutants G1092A, G1093A, G1092A G1093A, and the deletion construct Δdistal CC. The dotted line denotes the zero-current level. **c.** Group data for experiments in (b), shown as mean ± SEM (n = 5) or as individual experiments (open circles). Data were normalized to the peak current at +140 mV and 5.95 μM free Ca^2+^.

### Ca^2+^ binding leads to large conformational changes in rTRPM5

Our electrophysiological studies revealed that rTRPM5 has a high affinity for Ca^2+^ (EC_50_ ∼300 nM) (Extended Data Fig. 1b-e), and that channel activation is rapidly followed by full desensitization (Fig. 1a, Extended Data Fig. 1f-h). To attempt to capture the channel in a Ca^2+^- bound state, we determined the structure of rTRPM5 in a condition that contained trace Ca^2+^ (i.e., in a buffer containing no added Ca^2+^ or Ca^2+^ chelator, where free [Ca^2+^] is in the low micromolar range^33^). Under these conditions, the cryo-EM data revealed a large conformational ensemble.

Parsing of the data using the cryoSPARC 3D variability utility coupled with heterogeneous refinement^34,35^ allowed us to identify 3 distinct classes (rTRPM5_trace-1_, rTRPM5_trace-2,_ and rTRPM5_trace-3_), each with a unique conformation of the cytoplasmic domains (Extended Data Fig. 8), that were of high enough resolution to allow building of atomic models. All three were determined in C4 symmetry. While some density was present around the distal CC, we could not resolve the individual helices for this region in any of these 3D reconstructions, even without symmetry imposed. This is consistent with a high degree of flexibility in this region which is suggested by our electrophysiological recordings of GG mutants and deletion of the distal CC (Fig. 4). Only one of the maps (rTRPM5_trace-3_) possessed density where Ca^2+^ could be unambiguously placed in the two previously determined binding sites^11^(Extended Data Figs. 9 and 10). We did not model Ca^2+^ ions in the rTRPM5_trace-1_ and rTRPM5_trace-2_ maps because the density in the Ca^2+^ binding sites was weaker. However, given the high affinity of the channel for Ca^2+^, it is likely that the ion is bound in these structures but not visible due to insufficient local resolution. We also observed non-protein density within the putative PIP_2_ binding site in all three structures (Extended Data Fig. 11a-c).

Importantly, the 3 structures differ from each other by the distance between neighboring cytoplasmic protomers. We organized the structures according to the distance between the MHR1/2 of one protomer (Cα K94) and the rib helix of the neighboring protomer (Cα N1090), starting from the longest (rTRPM5_trace-1_, ∼22 Å) to the shortest (rTRPM5_trace-3_, ∼12 Å) (Fig. 5a, Extended Data Fig. 12a). By comparison, in the tightly coupled zebrafish TRPM5 this distance measures ∼ 11 Å^11^ (Extended Data Fig. 7e). Because the distance between protomers at this interface in rTRPM5_trace-3_ is similar to that of the zebrafish TRPM5, we will refer to rTRPM5_trace- 3_ as fully coupled.

**Fig. 5.**
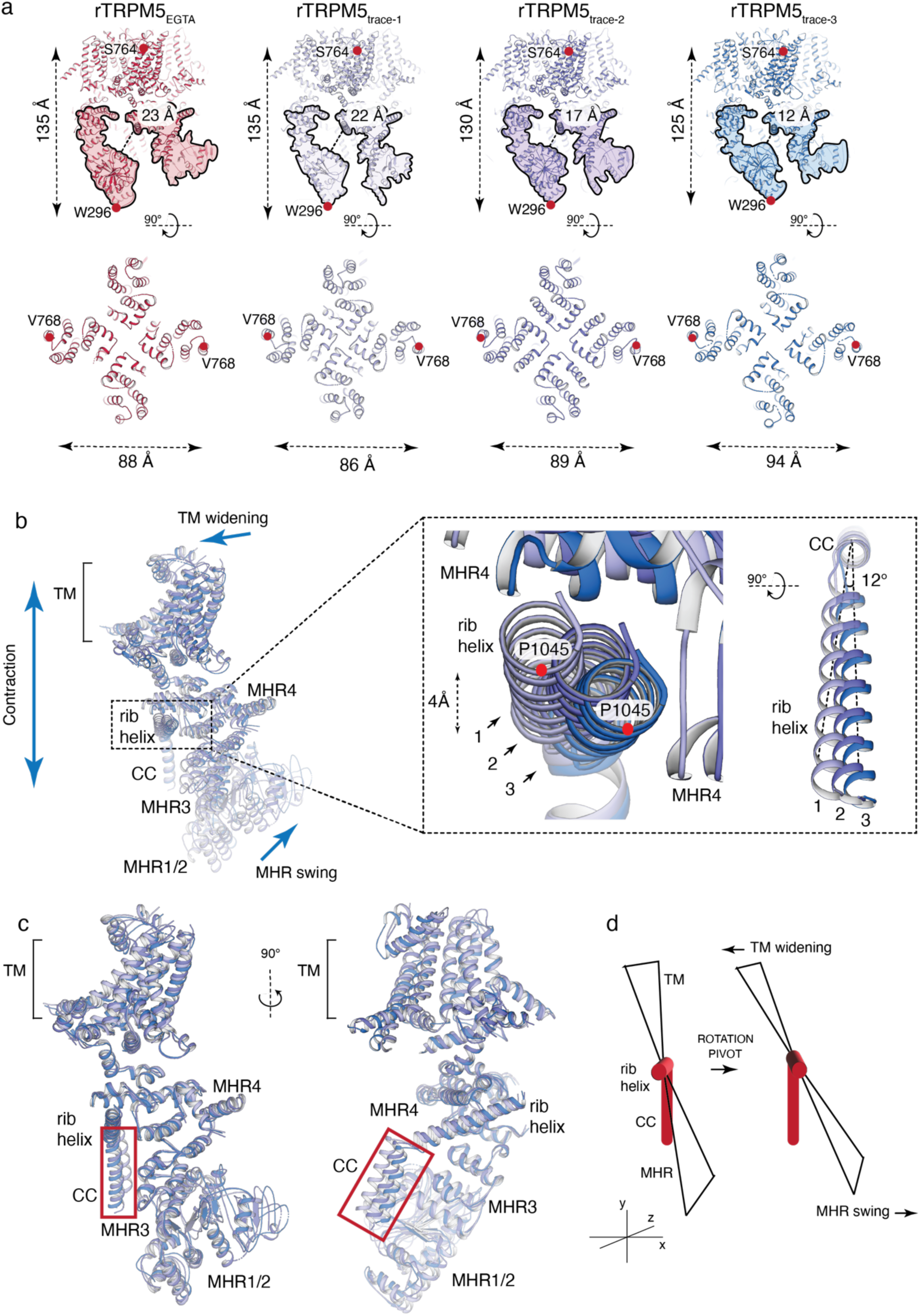
Trace amounts of calcium induce large conformational changes in rTRPM5. **a** Comparison of rTRPM5_EGTA_ (view of MHR A and D, as shown in red in Fig. 3a) with the conformations of rTRPM5 determined in the presence of trace amounts of Ca^2+^ (rTRPM5_trace-1_, lilac; rTRPM5_trace-2_, purple; rTRPM5_trace-3_, blue). Top panel shows side view. The cytoplasmic domains are highlighted. Dashed line represents the distance between MHR1/2 helix Cα K94 and the rib helix Cα N1090. The length of the channel (the distance between the Cα carbons of the S2 residue S764 and the MHR1/2 residue W296) condenses from 135 Å in rTRPM5_trace-1_ to 125 Å in rTRPM5_trace-3_. The bottom panel shows top view. The shortening of the channel coincides with a widening of the TM domains, as measured from the Cα of S2 residues V768 of opposing protomers. The TM domains widen from 86 Å in rTRPM5_trace-1_ to 94 Å in rTRPM5_trace-3._ **b** Alignment of rTRPM5_trace-1,_ rTRPM5_trace-2,_ and rTRPM5_trace-3_ tetramers. For clarity, only a single protomer is shown for each structure. The alignment reveals a rotation around the rib helix (indicated by box and enlarged inset). The rib helix of rTRPM5_trace-3_ is rotated by 12° and displaced laterally by ∼4 Å compared to rTRPM5_trace-1_. The red dot signifies the position of P1045 in the rib helix. The extent of the rib helix rotation correlates with the extent of channel shortening and TM widening described in **a**. **c** An overlay of the individual protomers shown in **b**. The agreement between the protomers is good across the protomer, except in the CC (red box). This indicates that the differences observed between rTRPM5_trace_ structures stem from the angle between CC and rib helix – i.e., rotation and pivot of the rib helix. **d** A schematic describing the conformational changes observed in rTRPM5_trace._

As observed earlier, as the interface between MHR domains becomes tighter, the length of the channel contracts: the rTRPM5_EGTA_ protomer A and rTRPM5_trace-1_ measure ∼ 135 Å from the top of the S2 (S764) to the bottom of MHR1/2 (W296). By contrast, the rTRPM5_trace-2_ is shortened to 130 Å and the rTRPM5_trace-3_ to 125 Å (Fig. 5a), similar to protomers C and D in rTRPM5_EGTA_ (Fig. 2b). However, in contrast with the C1-symmetric rTRPM5_EGTA_ structure, the movements in the cytoplasmic domains are accompanied by significant changes in the dimensions of the TM domain in the C4-symmetric rTRPM5_trace_ structures. Namely, as the channel contracts vertically, its TM domains expand laterally. When measured across the TM (S2, V768), the channel widens from 86 Å in rTRPM5_trace-1_, to 94 Å in rTRPM5_trace-3_ (Fig. 5a). The widening in the TM domain also coincides with a widening of the S6 gate of the channel (Extended Data Fig. 12b-c). Therefore, rTRPM5_trace-3_ – the only structure in the set with a coupled cytoplasmic interface – also has the widest opening at the S6 gate (Extended Data Fig. 12b-c). A comparison with the open conformation of the zebrafish TRPM5 channel^11^ reveals that the pore dimensions of rTRPM5_trace- 3_ are very similar to those of the open channel pore (Extended Data Fig. 12d). We will therefore refer to this structure as tentatively open.

To determine which conformational changes are required to transition between rTRPM5_trace-1_ and rTRPM5_trace-3_, we aligned the tetrameric channels (Fig. 5b). The overlay shows that in rTRPM5_trace-3_ the MHR domains swing upwards while the TM domains tilt downwards when compared to rTRPM5_trace-1_ and rTRPM5_trace-2_. Interestingly, we observe a 12° rotation and a 4 Å downward movement of the rib helix from rTRPM5_trace-1_ to rTRPM5_trace-3_ (Fig. 5b). To determine if this rotation is a part of a rigid body movement of the entire protomer, we superposed the protomers. This resulted in a near-perfect alignment (RMSD 1.5 Å), leaving only the proximal CC unaligned (Fig. 5c). This suggests the key difference between the protomers lies in the relationship between the proximal CC and the rest of the protomer. In other words, the entire protomer undergoes a rotation and pivot around the proximal CC, which acts as a stable, immobile stalk. Such a rotation and pivot can explain the widening of the TM domains and the contraction of the channel length (Fig. 5d).

It is interesting that we register no independent movement of MHR1/2 in the rTRPM5_trace_ structures. Instead, the entire protomer – including MHR1/2 – moves as a single rigid body. We speculate that Ca^2+^ binding at the interface between MHR1/2 and MHR3 stabilizes the MHR1/2 domain, and the concerted nature of that change is consistent the high degree of cooperativity for Ca^2+^ dependent activation of rTRPM5.

Given the high affinity of rTRPM5 for Ca^2+^, it is likely that the rTRPM5_trace_ ensemble represents desensitized states. However, desensitized channels still activate upon Ca^2+^ binding, although Ca^2+^-bound closed states are favored over open states after desensitization has occured^18,36^. This strong bias towards closed states after desensitization might explain why only rTRPM5_trace-3,_ with its coupled cytoplasmic interprotomer interfaces and open pore, was observed in a likely activated state. It has been a general observation that purified TRP channels in the presence of activators are rarely captured in the expected activated state in structural studies, suggesting that biochemical conditions generally tend to disfavor TRP channel opening^37^. It is important to note that, because density in the putative PIP_2_ site is present in all three structures (Extended Data Fig. 11a-c), our structures most likely reflect Ca^2+^-dependent desensitization of PIP_2_-bound channels. The contribution to desensitization of PIP_2_ depletion remains to be examined at the structural level. Mechanistically, these structures suggest that binding of Ca^2+^ stabilizes the MHR1/2 and induces a concerted rotation and pivot of protomers around the CC. These movements result in increased coupling between the protomers at the cytoplasmic interfaces, shortening of the channel along the y-axis, and widening of the pore.

### Rat TRPM5 adopts a range of desensitized, closed conformations in high Ca^2+^

Our electrophysiological data showed that prominent Ca^2+^ dependent desensitization is a hallmark of mammalian TRPM5 channels. To attempt to overcome the bias towards closed states post-desensitization and to ensure that Ca^2+^ ions can be unambiguously identified, we collected cryo-EM data of rTRPM5 in the presence of high (2 mM) Ca^2+^.

The 2 mM Ca^2+^ condition also yielded a large conformational ensemble. Using a similar approach to the one applied to the rTRPM5_trace_ dataset, we were able to identify 4 classes (rTRPM5_high-1_, rTRPM5_high-2,_ rTRPM5_high-3_, and rTRPM5_high-4_) which yielded 3D reconstructions of resolution sufficient for model building (Extended Data Fig. 13). Like in the rTRPM5_trace_ dataset, we could not resolve the individual helices corresponding to the distal CC in this dataset. Non-protein density was observed in the putative PIP_2_ site in all 4 classes (Extended Data Fig. 11d-g). In addition, density corresponding to Ca^2+^ was observed in both Ca^2+^ binding sites in rTRPM5_high-1_, rTRPM5_high-2,_ and rTRPM5_high-4_ (Extended Data Figs. 9 and 10). In rTRPM5_high-3_ we did not assign any Ca^2+^ densities. However, we speculate that – given the high rTRPM5 affinity for Ca^2+^ and the high Ca^2+^ concentration – the ion may be bound but is perhaps not fully visible due to local resolution.

The 4 structures obtained in 2 mM Ca^2+^ can be arranged according to the distance between protomers at the cytoplasmic interfaces. rTRPM5_high-1_ has the largest gap between MHR1/2 (Cα K94) of one protomer and the rib helix (Cα N1090) of the neighboring protomer (18 Å), and rTRPM5_high-4_ the smallest (∼12 Å) (Fig. 6a, Extended Data Fig. 14). Indeed, like in the rTRPM5_trace1-3_ structures, the protomers pivot and rotate around the CC to result in more compact length and wider TM domains (Fig. 6a, Extended Data Fig. 14a-b). However, the S6 gate in rTRPM5_high_ remains closed (Extended Data Figs. 14d and 15), and we therefore assume that all four conformations represent desensitized closed channels.

**Fig. 6.**
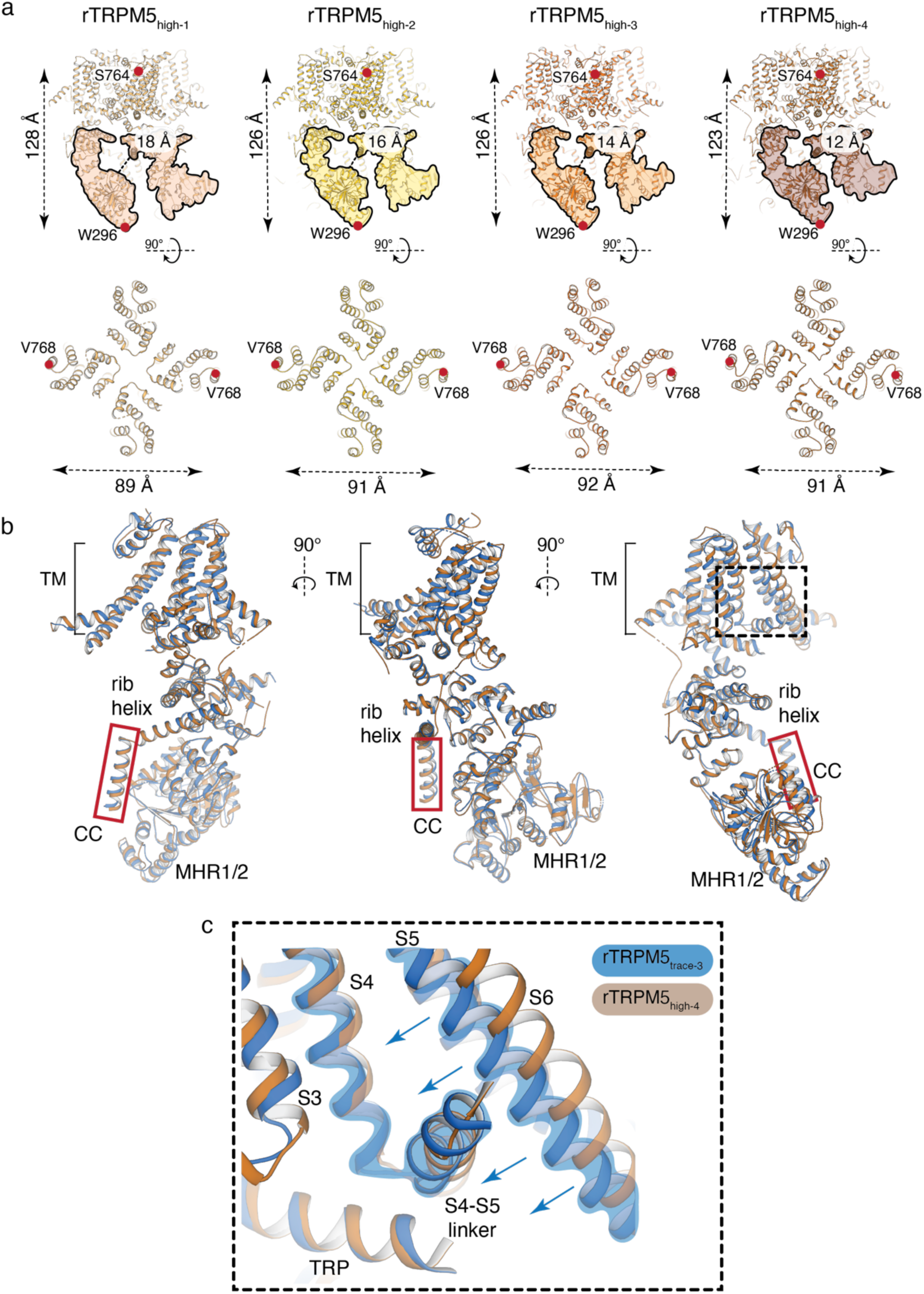
Structures of rTRPM5 in the presence of a high Ca^2+^ concentration. **a** A side by side comparison of rTRPM5_high1-4_ structures: rTRPM5_high-1_, light orange; rTRPM5_high-2_, gold; rTRPM5_high-3_, deep orange; rTRPM5_high-4_, maroon. Top panel shows a side view and the cytoplasmic domains are highlighted. Dashed line indicates the distance between MHR1/2 α2 helix Cα K94 and the rib helix Cα N1090 of neighboring protomers. The length of the channel is measured between the Cα of the S2 residue S764 and the MHR1/2 residue W296. Bottom panel shows top view. As observed in rTRPM5_trace_, the shortening coincides with widening of the TM domains. Notably, the differences in length/width dimensions are smaller within the rTRPM5_high_ structures than the rTRPM5_trace_ structures. **b** An overlay of individual protomers of rTRPM5_trace-3_ (blue) and rTRPM5_high-4_ (maroon) – structures which exhibit the highest degree of coupling in the two datasets. The protomers overlay well overall. Notable differences include a small offset in the position of the CC (shown in red box) and the conformation of the pore (dashed line box). **c** A close-up of the interface between S4, S4-S5 linker, S5, and S6 in rTRPM5_trace-3_ (blue) and rTRPM5_high-4_ (maroon). The S6 in rTRPM5_high-4_ is pulled by the S4-S5 linker to result in a wider conformation of the pore.

To interrogate the mechanisms of activation and desensitization in rTRPM5, we compared the tentatively open, fully coupled rTRPM5_trace-3_ with the closed, fully coupled rTRPM5_high-4_ (Fig. 6b). When single protomers of the two channels are superposed, we observe only a small change in the CC (Fig. 6b). This indicates that the extent of rotation and pivot around the rib helix is similar in the two channels, consistent with full Ca^2+^ occupancy in the two samples (Extended Data Figs. 9 and 10). Consequently, the extent of coupling between the MHR domains is also similar (∼ 12 Å distance between MHR1/2 and the rib helix of neighboring protomers, (Figs. 5a and 6a, Extended Data Figs. 12a and 14c). However, the overlay also shows that there are significant differences between the VSLD, the S4-S5 linker, and the pore domain in the two structures (Fig. 6b-c). In rTRPM5_trace-3_, the S4-S5 linker interacts with the S6 helix, which is positioned closer to the VSLD and away from the center of the pore, apparently pulling on the S6 gate to open the pore (Fig. 6c).

Our data suggests that four fully coupled cytoplasmic interprotomer interfaces are required to achieve opening of the rTRPM5 channel pore. Full coupling appears to require Ca^2+^ binding, as observed in rTRPM5_trace-3_ and rTRPM5_high-4_. By contrast, in the absence of Ca^2+^, e.g., in our rTRPM5_EGTA_ structure, the cytoplasmic interprotomer interfaces are highly dynamic, adopting a range of uncoupled and nearly coupled conformations (Fig. 2b-c). The remaining structures, obtained both under low and high Ca^2+^ conditions, have uncoupled cytoplasmic interprotomer interfaces and closed channel pores (Figs. 5a and 6a, Extended Data Figs. 12a, 14c, and 15), and likely represent desensitized closed states. We therefore posit that desensitization in rTRPM5 might be associated with the uncoupling of the cytoplasmic interprotomer interfaces in fully Ca^2+^- bound channels, which disengages critical networks that lead to gating. Desensitized channels still undergo gating and activate upon Ca^2+^-binding, but the gating equilibrium is biased towards closed states; the mechanistic and structural basis for this is presently unclear, but our data suggest it involves a weakening of the interactions at the cytoplasmic interprotomer interfaces.

### Coupling of cytoplasmic interfaces is necessary for rTRPM5 function

To interrogate the functional role of the cytoplasmic interprotomer interfaces and their significance in the activation and desensitization of rTRPM5, we took a closer look at the molecular interactions under uncoupled and coupled conditions.

None of the interprotomer interfaces in rTRPM5_EGTA_ are fully coupled (Fig. 2c). At the interface between protomers A and D the distance is largest – 23 Å (Fig. 2c, Fig. 7a). Here, the α2 helix of MHR1/2 has no interactions with the neighboring protomer (Fig. 7a-b). But positively charged residues, R89 and R85, in the α2 helix, extend towards the α3 helix within its own MHR1/2 and are positioned to make cation-pi contacts with H115 (Fig. 7a-b). These internal MHR1/2 interactions keep the α2 and α3 helices together. When the interface is coupled, e.g., as in rTRPM5_trace-3_ and rTRPM5_high-4,_ the α2 and α3 helices come within interaction distance to the rib helix and the MHR3 α4 helix of the neighboring protomer (Fig. 7c-d). Here, the internal interactions between helices α2 and α3 of MHR1/2 are preserved and appear critical for stabilizing the interface between α3 and the rib helix (Fig. 7d). At the interface between α2 and the neighboring MHR3 α4 helix, D404 in the MHR3 α4 is within interaction distance to several residues that include K81, S82, and R85 in MHR1/2 α2 (Fig. 7c-d). Based on our structural data, we hypothesized that the interfaces between MHR1/2 and rib helix and MHR3 of the neighboring protomer dictate the functional state of the channel and that a fully coupled interface is necessary for opening of the channel, both before and after desensitization.

**Fig. 7.**
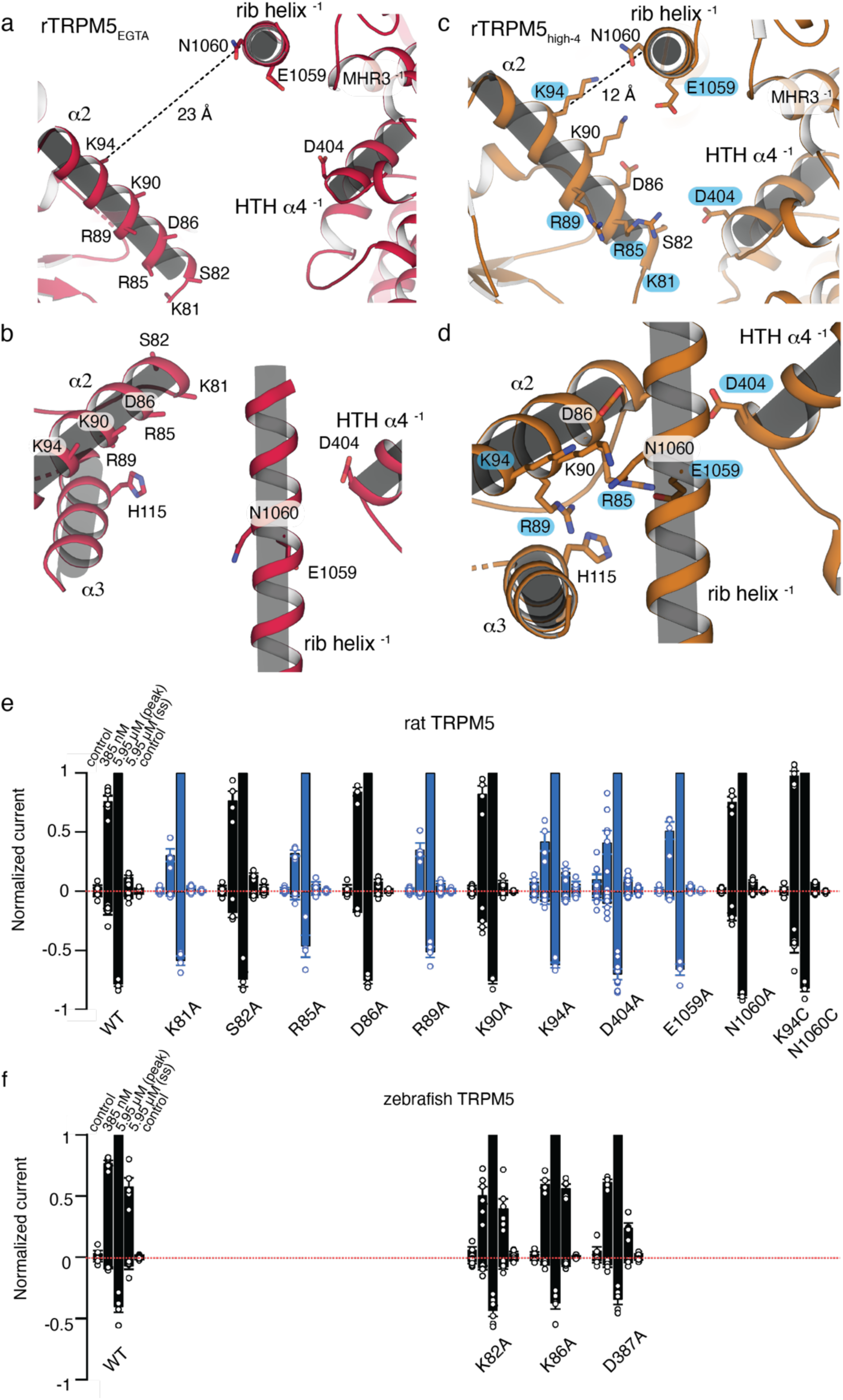
The interprotomer contacts between MHR1/2 and MHR3 and the rib helix are functionally important. **a-d** Interfaces between MHR1/2 and MHR3 and the rib helix of neighboring protomers in rTRPM5_EGTA_ (view of protomers A and D, as shown in red in Fig. 4a, a-b) and rTRPM5_high-4_ (c- d). **a** Side view of the interface between MHR1/2 and MHR3 and the rib helix from the neighboring protomer. Only MHR1/2 α2 helix is shown for clarity. The distance between the Cα atoms of K94 (MHR1/2) and N1060 (rib helix) measures ∼ 23 Å, indicating no interactions/coupling. **b** Top view of the interface in rTRPM5_EGTA_ including the α3 helix of the MHR1/2 domain. **c** Side view of the MHR1/2 and MHR3 and the rib helix interface in rTRPM5_high-4_. The distance between the Cα atoms of K94 and N1060 condenses to ∼ 12 Å. **d** Top view of the interface shows how α2 and α3 from MHR1/2 and HTH α4 of the neighboring MHR3 condense around the rib helix. The blue residue labels in c and d reflect results described in e. **e** Mean ± SEM inside-out patch currents from cells expressing WT or mutant rat TRPM5 channels as shown on Extended Data Figure 16. Data were measured at ± 140 mV, normalized to peak at +140 mV and 5.95 μM free Ca^2+^, and quantified at each of the indicated intracellular free Ca^2+^ conditions. Data from individual experiments are shown as circles (n = 4-8). Bars are colored black for WT and WT-like mutants, and blue for mutants with reduced responses to 385 nM free Ca^2+^. **f** Mean ± SEM inside-out patch currents from cells expressing WT- or mutant zebrafish TRPM5 channels as in Extended Data Fig. 16 and depicted as in **e** (n = 5-6).

To test this hypothesis, we introduced mutations into three distinct regions of this cytoplasmic interface: (1) the MHR1/2 α2 – rib helix inter-protomer interaction (mutations K90A, K94A, E1059A, N1060A, and K94C E1059C), (2) the MHR1/2 α2 – α3 intra-protomer interaction (mutations R85A, D86A, R89A), and (3) the MHR1/2 α2 – MHR3 α4 inter-protomer interaction (mutations K81A, S82A, and D404A). We recorded currents from inside-out patches under activating nanomolar concentrations of Ca^2+^ and under more strongly desensitizing Ca^2+^ conditions (Fig. 7e and Extended Data Fig. 16). Our data showed that disrupting the MHR1/2 α2 – MHR3 α4 interface by neutralizing D404 resulted in highly variable responses to nanomolar concentrations of Ca^2+^ (Fig. 7e and Extended Data Fig. 16). Mutations that disrupt the interactions between the two MHR1/2 helices, α2 and α3, also resulted in similar alterations of function. Namely, in mutants R85A and R89A we observed a significant decrease in current amplitude in response to nanomolar Ca^2+^ concentrations relative to their response to micromolar Ca^2+^ (Fig. 7e and Extended Data Fig. 16). The K94A and E1059A mutations, which disrupt the MHR1/2 α2 – rib helix interface, resulted in a similar phenotype (Fig. 7e and Extended Data Fig. 16a). Interestingly, other similar mutations at this interface, such as K90A, N1060A, D86A, and a double-cysteine mutant at the inter-protomer interface, K94C + N1060C, exhibit a WT phenotype (Fig. 7e and Extended Data Fig. 16a).

Together, these data are consistent with our hypothesis that the coupled conformation is necessary for proper rTRPM5 channel function. The effects of K81A and D404A suggest that the interface between MHR1/2 and MHR 3 of the neighboring protomer plays a role in rTRPM5 activation by Ca^2+^. It is interesting to note that this interface is small, created by interactions that form at the tips of two helices. It is therefore perhaps not surprising for a disruption to have an effect on channel activity. The intra-protomer interactions between the two MHR1/2 helices, α2 and α3, also appear to play a consequential role in channel activation, presumably because they help stabilize the interaction between α3 and the rib helix.

In contrast to rTRPM5, zebrafish TRPM5 channels containing mutation D387A, equivalent to D404A in the rat ortholog, exhibit a more WT-like behavior (Fig. 7f and Extended Data Fig. 16b). Similarly, mutations to the MHR1/2 – rib helix interface in the zebrafish TRPM5 had less marked effects on channel function (Fig. 7f). Taken together with the zebrafish TRPM5 structures, which show that these channels adopt a coupled conformation under a wide range of experimental conditions^11^ (Extended Data Fig. 7f), these results suggest that the interprotomer interface in the zebrafish channel is much more robust and less dynamic than that in rTRPM5, and therefore possibly less prone to desensitization. Indeed, in the zebrafish channel the cytoplasmic inter- protomer interface also involves significant interactions between MHR4 domains of neighboring protomers as well as MHR4 and the coupling domain^11^ (Extended Data Fig. 7). Consistent with this, point mutations at the interprotomer interfaces between MHR1/2 – MHR3 and MHR1/2 – rib helix have a minimal effect on zebrafish channel activation and calcium sensitivity.

To summarize, we found that the rat TRPM5 channel exhibits a large degree of conformational flexibility in the presence of Ca^2+^ and is sensitive to mutations that destabilize the cytoplasmic interprotomer interface. In stark contrast, the zebrafish ortholog has markedly reduced desensitization, and rigid cytoplasmic interprotomer interfaces. Our structural and functional data suggest that full coupling at interfaces formed by MHR1/2 – MHR3 and MHR1/2 – rib helix is necessary for activation of rTRPM5 by intracellular Ca^2+^ but also that interactions at these interfaces become weakened both in the absence of stimulus and upon desensitization. This might therefore mean that the labile cytoplasmic interfaces in rTRPM5 are a critical feature that endow the channel with enhanced flexibility and thus an extended range of possible modulatory mechanisms. The zebrafish channels do not share the requirement for desensitization, and this may be reflected in their robust interprotomer interfaces.

### Both Ca^2+^ binding sites contribute to rTRPM5 activation

To determine whether channel activation or desensitization might be driven preferentially by one of the two Ca^2+^ binding sites, we mutated both the VSLD site (Extended Data Fig. 17a) and the ICD site (Extended Data Fig. 17b) and tested their response to activating nanomolar Ca^2+^ concentrations and desensitizing micromolar Ca^2+^ concentrations. Mutating residue Q784 in the VSLD binding site, which in the zebrafish channel had proven critical for channel activation^11^, resulted in channels that lacked a response to nanomolar Ca^2+^ concentrations, and exhibited currents of reduced magnitude in response to micromolar Ca^2+^ relative to WT channels (Extended Data Fig. 17c). In the ICD site, the single mutation E354A had a slightly reduced response to nanomolar Ca^2+^ concentrations but responded to micromolar Ca^2+^ similarly to WT channels (Extended Data Fig. 17c). However, more invasive disruptions of the ICD Ca^2+^ site via double, triple, quadruple, and total ablation of the site (i.e., a quintuple mutation), dramatically reduced the rTRPM5 channel’s ability to respond to Ca^2+^ to the extent that currents we recorded from patches expressing these mutants were indistinguishable from endogenous currents recorded from GFP-transfected control cells (Extended Data Fig. 17c). Our data indicate that both Ca^2+^ binding sites are essential for Ca^2+^-dependent activation, because disruption of each site resulted in apparent loss of TRPM5 channel function.

It is also interesting to note that the outcomes of mutations to the VSLD and ICD Ca^2+^ binding sites appear to differ between the zebrafish and rat TRPM5 channels. For example, mutation Q784A at the VSLD site (rat TRPM5 numbering) abolishes the zebrafish TRPM5 current^11^ but rat TRPM5 mutants can still produce robust currents in response to micromolar Ca^2+^. Similarly, E354A mutation in the ICD site of the zebrafish channel renders the channel more voltage sensitive at high Ca^2+^ concentrations^11^, whereas the rat mutant channel behaves very similarly to WT (Extended Data Fig. 17c). These observations point to subtle differences in how the Ca^2+^-mediated response is choreographed in the two orthologs and invite future studies to dissect the intricacies of this system.

## Discussion

Here we have described the conformational changes that occur during gating in a mammalian TRPM5 channel (Fig. 8a). In the absence of Ca^2+^, the cytoplasmic assembly is highly flexible and its dynamics appear to be uncoupled from the transmembrane domain and pore. The cytoplasmic interfaces between protomers adopt a range of uncoupled conformations, apparently independently of each other. The binding of Ca^2+^ sets in motion a series of concerted transitions, reflected in the extremely high cooperativity of activation by Ca^2+^, which couple the movements at the cytoplasmic assembly with the conformation of the transmembrane domain and pore. Once Ca^2+^ binds to all 8 sites, each protomer undergoes rotation and pivot around the CC, and the MHR1/2 is re-positioned to engage with the neighboring protomer. Once the channel becomes fully coupled the pore gate can open. Under desensitizing conditions, the interactions at the cytoplasmic interfaces are likely weakened, contributing to the range of distances between protomers at the cytoplasmic interface that we observe in all structures determined in the presence of Ca^2+^. We therefore posit that in mammalian TRPM5 channels the tight coupling at the cytoplasmic interfaces occurs only in the active state. During desensitization, the tight coupling is destabilized and replaced by enhanced dynamics of the cytoplasmic interface.

**Fig. 8.**
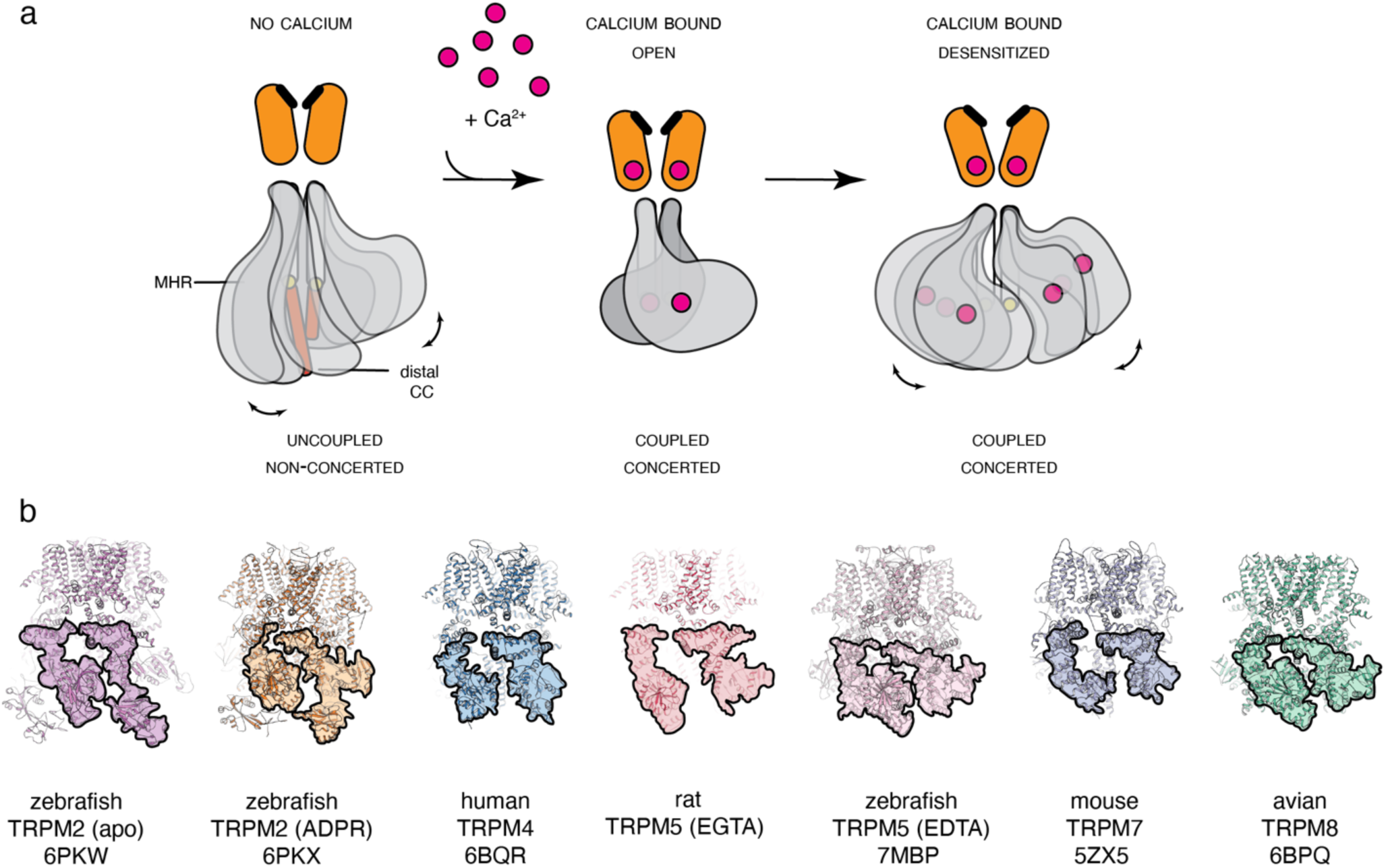
The role of cytoplasmic domains in mammalian TRPM function. **a** A cartoon representation of the proposed rTRPM5 mechanism. In the absence of Ca^2+^, the cytoplasmic domains possess a high degree of flexibility and can be observed in a range of conformations. However, the cytoplasmic interprotomer interfaces remain uncoupled. The channels can open when all Ca^2+^ sites are fully occupied and the cytoplasmic interprotomer interfaces are fully coupled. Finally, during desensitization a relaxation occurs - the coupling between protomers decreases, the MHR domains become flexible again, and the open state becomes destabilized. Importantly, conformational changes remain concerted between all four subunits in the presence of Ca^2+^. **b** Cytoplasmic interfaces in some existing TRPM structures. Shown are zebrafish TRPM2 (6PKW and 6PKX), human TRPM4 (6BQR), rat TRPM5 in EGTA (8SL6, this study), zebrafish TRPM5 in EDTA (7MBP), mouse TRPM7 (5ZX5) and avian TRPM8 (6BPQ). Cytoplasmic domains have been shaded for clarity.

A fascinating aspect of the mechanism of mammalian TRPM5 channel modulation by Ca^2+^ is that its binding triggers both activation and desensitization. We found that disruption of either of the two identified Ca^2+^ sites is sufficient to prevent channel activation, suggesting that occupancy at both sites is required for activation. This is further supported by the steep cooperativity associated with channel activation by Ca^2+^. It remains to be determined whether any additional binding sites exist that mediate entry into the desensitized state. The processes of desensitization and activation, however, seem to have a very similar dependence on Ca^2+^- concentration. We therefore hypothesize that desensitization requires occupancy of the same sites as those that mediate activation; initial Ca^2+^-binding promotes occupancy of a non-desensitized activated state, which rapidly transitions into a kinetically absorbing ensemble of desensitized states where channel opening is energetically disfavored. The role of PIP_2_ in this process also requires additional investigation, but our results establish that increasing the amount of the soluble analogue diC8 PIP_2_ can almost entirely restore channel activity to pre-desensitization levels at high Ca^2+^ concentrations. This had been previously observed in the closely related TRPM4 channel, albeit at lower diC8 PIP_2_ concentrations^38^. All of our structures contain densities that are consistent with PIP_2_ at a site that is conserved in TRPM8^31^ and TRPM3 channels^39^ and further studies will be needed to determine the mechanism underlying PIP_2_-dependent desensitization of rat TRPM5. This study confirms the functional similarities between the TRPM4 and TRPM5 channels, such as Ca^2+^-dependent desensitization and PIP_2_ dependence. The study also draws attention to a shared importance of the cytosolic interprotomer interface in the function of the two channels.

Namely, in TRPM4 this interface contains the binding site for ATP, an inhibitor of TRPM4 function, as well as decavanadate, a channel modulator^28^. ATP apparently inhibits the channel by uncoupling of the interprotomer interface^29^. Moreover, several of the TRPM4 mutations that have been identified in patients with inherited cardiac disease occur at the interface between MHR1/2 and MHR3 of neighboring protomers^40,41^.

Interestingly, coupling and uncoupling at the cytoplasmic interprotomer interfaces also occurs during gating of the more distantly related TRPM2^42,43^. This hints at the possibility that the cytoplasmic interfaces could be important for gating in several types of TRPM channels. Indeed, a cursory look at the cytoplasmic interfaces in the available TRPM channel structures (a small selection is shown in Fig. 8b) reveals many different arrangements of the cytoplasmic domains. This presents us with the question: wwhat do these distinct arrangements tell us about their modes of gating? Additional studies of TRPM conformational ensembles might enable us to answer this question and understand where TRPM gating mechanisms intersect and where they diverge.

## Methods

### Protein Expression and Purification

A codon optimized gene for rat (*Rattus norvegicus*) TRPM5 was synthesized and cloned into a customized pEG BacMam vector in frame with an N-terminal FLAG affinity tag. Rat TRPM5 plasmid DNA transformed into DH10Bac *E. coli* cells and baculovirus was obtained as previously described^44^. 5-10% (v/v) of P3 baculovirus was added to HEK293 GnTI^-^ cells at densities of ∼2.5-3 M/mL in Freestyle 293 Media supplemented with 2% (v/v) FBS and 1% (v/v) penicillin-streptomycin. Cultures were grown at 37°C, 120 rpm, and 8% CO_2_ for ∼18 hours after which 10 mM sodium butyrate was added and the temperature was lowered to 30°C. After an additional ∼48 hours, cells were pelleted and resuspended in a buffer A (150 mM NaCl, 50 mM Tris pH 8.0, 2 mM DTT) supplemented with a protease inhibitor cocktail (1 μg μL^-1^ leupeptin, 1.5 μg μL^-1^ Pepstatin, 0.84 μg μL^-1^ aprotinin, 0.3 mM PMSF), 14.3 mM BME, DNAse I, and 2 mM DTT. For the calcium-free EGTA preparation, 1 mM EGTA was also included.

Cells were solubilized for 1 hour at 4 °C in 1% (w/v) Glyco-diosgenin (GDN) (Anatrace). Insoluble material was removed by centrifugation at 16,000 rpm for 30 minutes at 4 °C. The supernatant was then incubated with anti-FLAG M2 resin (Millipore-Sigma) and allowed to incubate for one hour at 4 °C. The supernatant – resin suspension was poured onto a Bio-Rad Econo-Column, and the flow-through was discarded. The remaining resin was subjected to 10 column volume (CV) washes with Buffer B (150 mM NaCl, 50 mM Tris pH 8.0, 2 mM DTT, 0.04% GDN, and 1 mM EGTA where noted). Elution was performed with 5 CV of Buffer C (150 mM NaCl, 50 mM Tris pH 8.0, 2 mM DTT, 0.04% GDN, 0.1 mg mL^-1^ FLAG peptide, and 1 mM EGTA where noted).

The eluate was concentrated and loaded onto a Superose 6 column (Cytiva) equilibrated with Buffer B. Peak fractions were concentrated to ∼ 3-4 mg mL^-1^ using a 100 kDa MWCO centrifugal concentrator (Millipore Sigma) for cryo-EM sample preparation.

### Cryo-EM sample preparation

For the rTRPM5_EGTA_ preparation, the sample was prepared in buffer B supplemented with 1 mM EGTA. The rTRPM5_trace_ and the rTRPM5_high_ samples were prepared in buffer B. After concentration, the rTRPM5_high_ sample was supplemented with 2 mM CaCl_2_ and incubated on ice for 30 minutes. Before vitrification, all samples were centrifuged for 30 minutes at 4 °C and 16,000 rpm on a tabletop micro-centrifuge to remove any precipitation.

A similar vitrification protocol was used for each preparation. For the rTRPM5_trace_ and rTRPM5_high_ samples, 3 μl sample was dispensed on a freshly glow discharged (30 s) UltrAuFoil R1.2/1.3 300- mesh grid (Electron Microscopy Services), blotted for 1-3 s with Whatman no. 1 filter paper using the Leica EM GP2 Automatic Plunge Freezer at 23° C and > 85% humidity and plunge-frozen in liquid ethane cooled by liquid nitrogen. For the calcium-free sample, the same protocol was followed but the protein was vitrified onto Quantifoil R1.2/1.3 300-mesh grid (Electron Microscopy Services).

### Cryo-EM data collection

Data for calcium-free, trace-calcium, and high calcium conditions were collected using the 300 keV Titan Krios transmission electron microscope (TEM) with either Gatan K3 or Falcon III Direct Electron Detectors operating in dose counting mode in conjunction with a Bioquantum K3 Imaging Filter operating at 20eV. Nominal magnifications used for the acquisitions were 81,000x corresponding to physical pixel sizes of 1.08 Å/pixel (0.54 Å/pixel super-resolution), 1.08 Å/pixel,

1.08 Å/pixel, and 1.12 Å/pixel for calcium-free, trace-calcium, and high calcium, respectively. For the calcium-free TRPM5 sample, 4,778 movies (40 frames/movie) were collected using an exposure rate of ∼14.0 e^-^/pixel/s for ∼2.3 s resulting in a total nominal exposure of 40 e^-^/Å. The target defocus range for each dataset was from −1.0 µm to −2.25 µm. For the trace calcium TRPM5 sample, 6,565 movies were collected (40 frames/movie) with 3.2 s exposure and exposure rate of ∼15.8 e^-^/pixel/s. The total exposure was of ∼45 e^-^/Å and a nominal defocus range from −1.0 µm to −2.5 µm. For the high calcium TRPM5 sample, 10,055 movies (40 frames/movie) were collected using an exposure rate of ∼16.7 e^-^/pixel/s for 2.8 s exposure resulting in a total nominal exposure of ∼43.3 e^-^/Å and had a nominal defocus range from -1.0 µm to -2.25 µm.

### Cryo-EM data processing rTRPM5_EGTA_

For TRPM5 in 1mM EGTA, all data processing was conducted using cryoSPARC V3.3.2^45^. Patch motion correction^46^ was used on the 4,778 dose-weighted movies. The motion corrected micrographs were then subject to patch CTF estimation^45^ after which they were curated by removal of those where total full-frame motion was > 40 pixels, estimated ice thickness was > 1.1, and resolution estimations were > 5 Å. This yielded a stack of 3,318 movies. From these we picked a total of 1,462,359 particles using the template picker (160 Å particle-size; 6 Å lowpass filter). The particles were extracted, binned at 4x4 (2.16 Å/pixel, 180 pixel box size), and subjected to 2D classification, which yielded 180,560 good particles. These particles were then re-extracted to 2x bin (1.08 Å/pixel, 360 pixel box size) and subjected to 5-class *ab initio* reconstruction in both C1 and C4 symmetries. From there, particles that contributed towards the best TRPM5 reconstructions (159,140 particles, C1; 172,214 particles, C4) were subjected to Non-uniform refinement^47^ that resulted in ∼4.2 Å and ∼3.8 Å volumes for C1 and C4 symmetries, respectively. However, the C1 imposed map had a greater coverage in the cytoplasmic domains (and especially the coiled coil domains) and the C4 maps were therefore not pursued further. The C1 map was used as input into 3D variability analysis^35^ (simple, 3-mode, 5 Å filter). Using 3D variability display, the derived volumes that showed the greatest articulations in the MHR domains were used to bin the original particle set using cryoSPARC heterogeneous refinement. The highest resolution heterogeneous volumes and particles were then subjected to another round of Non-uniform refinement (86,152 particles) which resulted in a 4.32 Å map in C1. This C1 map was used for model building due to its increased coverage of the distal coiled coil region.

### rTRPM5_trace_

All data processing was conducted using cryoSPARC V3.3.2. Patch motion correction utility was used on the 6,565 dose-weighted movies. The motion corrected images were then subject to patch CTF estimation after which they were pruned for total full-frame motion >40 pixels, estimated ice thickness > 1.1, and CTF estimations of > 5 Å. Using a combination of manual and template-based picking in conjunction with reference-free 2D classification, a set of 149,532 particles was selected to build a 1-Class *ab initio* reconstruction with C4 symmetry imposed. These particles were subjected to non-uniform refinement (4.12 Å), which was used to build a 50-class 2D template for picking. Using the optimal template (160 Å particle-size; 6 Å lowpass filter), 2,601,822 particles were picked from 6,042 movies. These particles were extracted, binned at 2x2 (2.16 Å/pixel, 180 pixel box size) and subjected to multiple rounds of reference-free 2D classification to identify 546,821 good particles. These particles were then re-extracted to full resolution (1.08 Å/pixel, 360 pixel box size) and subjected to 5-class *ab initio* reconstruction with C4 symmetry. From there, particles that contributed towards good TRPM5 reconstructions (510,640 particles) were subjected to a second round of 5-class *ab initio* (475,743 particles, C4) which was used to further curate the set. These particles were input into a non-uniform refinement (3.74 Å) in C4. These particles were then symmetry expanded to C4 and used as input into 3D variability analysis (simple, 3-mode, 3.5 Å filter). Using 3D variability display, the derived volumes that showed the greatest articulations in the MHR domains were used as reference volumes in heterogeneous refinement with C4 symmetry imposed. The highest resolution heterogeneous volumes and particles were then subjected another round of non-uniform refinements with the best particle and volume combinations produced 6.02 Å (49,620 particles), 3.81 Å (139,580 particles), and 4.17 Å (53,230 particles) maps. These particles were C4 symmetry expanded and subject to local refinements with a mask covering either the MHR and the coiled-coil domain of a single protomer or the full-length (FL) protein. The resulting full-length protein maps were 4.31Å, 3.70 Å, and 4.19 Å, while their respective focused maps of the MHR domains were 5.12 Å, 4.03 Å, and 4.40 Å.

### r TRPM5_high_

For TRPM5 in 2 mM Ca^2+^, all data processing was conducted using cryoSPARC V3.3.2. Patch motion correction utility was used on the 10,055 dose-weighted movies. The motion corrected images were then subject to patch CTF estimation after which they were pruned for total full-frame motion > 40 pixels, estimated ice thickness > 1.1, and CTF estimations of > 5 Å. Using the optimal template (160 Å particle-size; 10 Å lowpass filter) derived from trace Ca^2+^, the remaining 7,003 movies were picked resulting in 5,766,941 particles. These particles were then extracted, 2x2 binned (2.16 Å/pixel, 180 pixel box size), and subjected to multiple rounds of cryoSPARC reference-free 2D classification to identify 512,251 good particles. These particles were then re-extracted to full resolution (1.08 Å/pixel, 360 pixel box size) and subjected to 2-class *ab initio* reconstruction with C4 symmetry. From there, particles that contributed towards good TRPM5 *ab initio* reconstructions (377,692 particles) along with the best class volume were input to non-uniform refinement to generate a 3.72 Å map in C4 symmetry. A second round of 5-class *ab initio* was used to further curate good particles (331,548 particles) which were then input into another non-uniform refinement that resulted in a 3.65 Å map in C4. These particles were then symmetry expanded to C4 and used as input into 3D variability analysis (simple, 3-mode, 4.0 Å filter). Using 3D variability display, the derived volumes that showed the greatest articulations in the MHR domains were then used as reference volumes in the cryoSPARC heterogeneous refinement in C4. The highest resolution heterogeneous volumes and particles were then subjected another round of non-uniform refinement with the best particle and volume combinations producing maps of 4.15 Å (44,520particles), 4.81 Å (36,043 particles), 3.87 Å (41,013 particles), and 4.52 Å (29,977 particles). These particles were C4 symmetry expanded, subject to local refinements with a mask of either the MHR and the coiled-coil domain of a single protomer or of the full-length (FL) protein. The resultant full-length protein maps from local refinements were 4.15Å, 4.49 Å, 3.85 Å, and 4.90 Å, while their respective MHR maps were 3.68 Å, 3.98 Å, 3.50 Å, and 4.22 Å.

All resolution estimates were based on the gold-standard FSC 0.143 criterion as calculated by cryoSPARC^48,49^.

### Model building

The rTRPM5_EGTA_ model was built directly into the cryo-EM density using the previously determined structure of the zebrafish TRPM5 (PDB ID 7MBP) as a starting reference. The subsequent models were built using the rTRPM5_EGTA_ as a starting reference. All models were built in Coot 0.98^50^. In rTRPM5_trace_ and rTRPM5_high_ datasets, the building of MHR domains was guided by the focused maps of these regions. The Molprobity^51^ server was used to guide the refinement. At the final stage, models were subjected to a round of real space refinement using the PHENIX 1.20 *real space refine*^52^ utility. Analysis and illustrations were performed using Pymol^53^ and UCSF Chimera^54^.

### Free Ca^2+^ concentration measurements with Fura-2

A Fura-2 pentapotassium salt (Invitrogen) stock was prepared at 1 mM in water. Standard calcium solutions were prepared in triplicate using the Calcium Calibration Buffer Kit #1 (Molecular Probes Cat. No. C3008MP), following the manufacturer’s instructions, and supplemented with Fura-2 to a final concentration of 50 μM. 120 μL of each standard solution were loaded on an opaque bottom 96-well plate, and read on a SpectraMax M3 fluorescence plate reader (Molecular Devices) using SoftMax Pro software (Molecular Devices). Excitation spectra were measured from 250 to 450 nm with an increment of 10 nm, and with emission at 510 nm. Calibration curves were constructed by plotting the logarithm of the free Ca^2+^ concentration for each standard solution vs log[(R-R_min_)/(R_max_-R) x (F^380^_max_/F^380^_min_)] (see Extended data figure 1a), where R is the ratio of 510 nm emission intensity with excitation at 340 nm, to 510 nm emission intensity with excitation at 380 nm; R_min_ is the ratio at zero free Ca^2+^, and R_max_ is the ratio at saturating Ca^2+^ (39 μM); F^380^_max_ is the fluorescence intensity with excitation at 380 nm for zero free Ca^2+^; F^380^_min_ is the fluorescence intensity with excitation at 380 nm for saturating free Ca^2+^.

We initially calculated the free Ca^2+^ concentrations in our experimental solutions using the MaxChelator software: https://somapp.ucdmc.ucdavis.edu/pharmacology/bers/maxchelator/CaEGTA-TS.htm.

Our solutions were composed of 150 mM NaCl, 10 mM HEPES, 5 mM EGTA and had a pH of 7.4 adjusted with NaOH. Using MaxChelator, we supplemented this solution with different concentrations of total calcium (using a 1M CaCl_2_ stock solution) to obtain desired concentrations of free Ca^2+^. We then used Fura-2 to measure the actual concentration of free Ca^2+^ in our recording solutions. First, we diluted our experimental solution samples at a ratio of 1:5 using a solution composed of 150 mM NaCl and 10 mM HEPES (pH 7.4) and supplemented with Fura-2 such that its final concentration after mixing both solutions would be 50 μM. This had two advantages: first, we avoided variations in the amount of Fura-2 between samples, and second, all fluorescence readouts fell within the optimal range of the calibration assay. For our experimental solution containing a higher calculated concentration of free Ca^2+^ of 1 μM, we prepared dilutions at 1:5, 1:20 and 1:50. Notably, the measured Ca^2+^ concentration after adjusting for each of the dilution factors was similar in all samples, indicating that our dilution procedure was unlikely to affect results for the rest of the samples. All solution samples were prepared in triplicate and measured together with a 1:5 dilution of our recording solution without any added calcium (zero Ca^2+^ condition), and a 1:5 dilution of our recording solution supplemented with enough calcium to reach a calculated free Ca^2+^ concentration of 2 mM (saturating Ca^2+^ solution). We measured the excitation spectra for the samples as done for the standard solutions, computed log[(R-R_min_)/(R_max_- R) x (F^380^ /F^380^)] using the zero-Ca^2+^ and saturating Ca^2+^ sample dilutions described above, and plotted these values against the logarithm of the free Ca^2+^ concentration, starting with the calculated theoretical values divided by 5 (see Extended data figure 1a). We obtained the free Ca^2+^ concentration for each of our samples by shifting its x-axis value such that the data would align with our calibration standard curve. On average, the measured free Ca^2+^ concentrations were ∼ 4.5-fold larger than the calculated concentrations. We used this value to estimate the concentrations of solutions containing 5 and 50 μM theoretical free Ca^2+^ concentrations. For solutions with calculated concentrations > 50 μM, we did not adjust concentrations.

### Cell culture for patch-clamp electrophysiology

HEK293 cells were kept at 37 °C with 5% CO_2_ and grown in Dulbecco’s modified Eagle’s medium (DMEM) with high glucose, pyruvate, L-glutamine, and phenol red, supplemented with 10% fetal bovine serum (vol/vol) and 10 mg ml^-1^ gentamicin. For transfection, cells were detached with trypsin, re-suspended in DMEM and seeded onto 3 mL dishes with No. 1 glass coverslips that were pre-treated for 30 minutes with Fibronectin (0.1 mg ml^-1^ in PBS, Millipore Sigma). Transfections were performed on the same day using FuGENE6 HD Transfection Reagent (Roche Applied Science). Channel constructs were co-transfected with pGreen-Lantern (referred to as GFP in the text, Invitrogen) at a ratio of 1:1. Patch-clamp recordings were done 18-36 h after transfection.

### Patch-clamp electrophysiology

Patch clamp recordings were performed on transiently transfected HEK293 cells at room temperature (21–23°C) using the inside-out configuration. Data were acquired and digitized with a dPatch amplifier system and SutterPatch software (Sutter Instrument, Novato CA), and analyzed using Igor Pro 8.04 (Wavemetrics, Portland, OR). Pipettes were pulled from borosilicate glass (1.5 mm O.D. x 0.86 mm I.D. x 75 mm L; Harvard Apparatus) using a Sutter P-97 puller and heat-polished to final resistances between 1 and 5 MΩ using a MF-200 microforge (World Precision Instruments). An agar bridge (1M KCl; 4% weight/vol agar; teflon tubing) was used to connect the ground electrode chamber and the main recording chamber. The intracellular and pipette solution consisted of (mM): 150 NaCl, 10 HEPES, 5 EGTA, pH 7.4 (NaOH). The diC8 PIP_2_ (Echelon Biosciences, Salt Lake City, UT) stock solutions were made in water at 1 mM (for 20 μM final concentration) or at 10 mM (for 200 μM final concentration). For 385 nM and 5.95 μM free Ca^2+^, the total concentration of calcium added to the recording solution was 2.23 and 4.45 mM, respectively.

A gravity-fed motorized perfusion system (RSC-200, BioLogic, France) was used to switch between recording solutions. Currents were elicited by ±140 mV pulses of 50 ms duration applied at 10 Hz from a holding potential of 0 mV. Data was acquired at 5 kHz and low-pass filtered at 1 kHz. The current time-courses shown on figures display the current average for the last 5 ms of each pulse, and were normalized to the peak current value at 5.95 μM free Ca^2+^ and +140 mV. For dose-response relations, the mean current in zero-Ca^2+^ control solution was subtracted from the current values at the various free Ca^2+^ concentrations, which were then normalized to the current at +140 mV in 552 nM (before desensitization) or 636 nM free Ca^2+^ (after desensitization). Curves were fit to the Hill equation, and for some experiments the Hill coefficient was fixed to a value of 8.

### Cell lines

HEK293GnTI^-^ cells were purchased from ATCC with authentication records. Additional authentication was not performed prior to this study and these cells were not tested for mycoplasma contamination. HEK293 cells from ATCC (CRL-1573) used for electrophysiology tested negative for mycoplasma.

### cDNAs used in structural and functional studies

We used the codon optimized gene for rat (*Rattus norvegicus*) TRPM5 cloned into a customized pEG BacMam vector in frame with an N-terminal FLAG affinity tag. Mouse TRPM5 in pcDNA3.1 was purchased from Addgene (plasmid #85189), human TRPM5 in pcDNA3.1+ (Accession No: NM_014555.3) was purchased from GenScript, and zebrafish TRPM5 in pEG BacMam was kindly provided by Seok-Yong Lee (Duke University).

## Data availability

The atomic coordinates and density maps have been deposited to PDB and EMDB under the following entries: rTRPM5_EGTA_ (8SL6, EMD-40574), rTRPM5_trace-1_ (8SL8, EMD-40575, EMD-40600), rTRPM5_trace-2_ (8SLA, EMD-40576, EMD-40599), rTRPM5_trace-3_ (8SLE, EMD- 40577, EMD-40597), rTRPM5_high-1_ (8SLI, EMD-40578, EMD-40596), rTRPM5_high-2_ (8SLP, EMD-40579, EMD-40595), rTRPM5_high-3_ (8SLQ, EMD-40580, EMD-40593), and rTRPM5_high-4_ (8SLW, EMD-40581, EMD-40592). Source data are available for this paper. All materials are available upon reasonable request.

## Acknowledgements

The initial biochemical optimization and vitrification of rTRPM5 was performed at the laboratory of Professor Seok-Yong Lee at Duke University. We are also thankful to Professor Lee for the kind gift of rat and zebrafish TRPM5 plasmids. Data were collected at National Cryo-Electron Microscopy Facility at the Frederick National Laboratory (NCEF) and at the Pacific Northwest Cryo-EM Center (PNCC). We thank Adam Wier, Ulrich Baxa (NCEF) and Omar Davulcu (PNCC) for assistance with cryo-EM data collection. We also appreciate the help of Zubcevic lab rotation students Alexandra Berkowitz, Peyton Oden, and Omar Tinoco at various stages of this project. This work was supported by start-up funds from The University of Kansas Medical Center and the Klingenstein-Simons Neuroscience Fellowship (to L.Z) and startup funds from the University of Texas at Austin (to A. J.-O.). This research was, in part, supported by the National Cancer Institute’s National Cryo-EM Facility at the Frederick National Laboratory for Cancer Research under contract 75N91019D00024. A portion of this research was supported by NIH grant U24GM129547 and performed at the PNCC at OHSU and accessed through EMSL (grid.436923.9), a DOE Office of Science User Facility sponsored by the Office of Biological and Environmental Research.

## Author Contributions

S.K and L.Z. prepared cryo-EM samples. S.K. performed all molecular biology. L.Z, S.K., and L.G.S. screened grids. L.G.S. performed cryo-EM data processing. L.Z., S.K., and L.G.S. built the atomic models and analyzed the structural data. A. J.-O. performed all electrophysiology experiments and analyzed the data. All authors discussed the results and commented on the manuscript.

## Competing Interests

The authors declare no competing financial interests. Correspondence and requests for materials should be addressed to L.Z. or A. J.-O.

## Extended Data Figures

**Extended Data Fig. 1.**
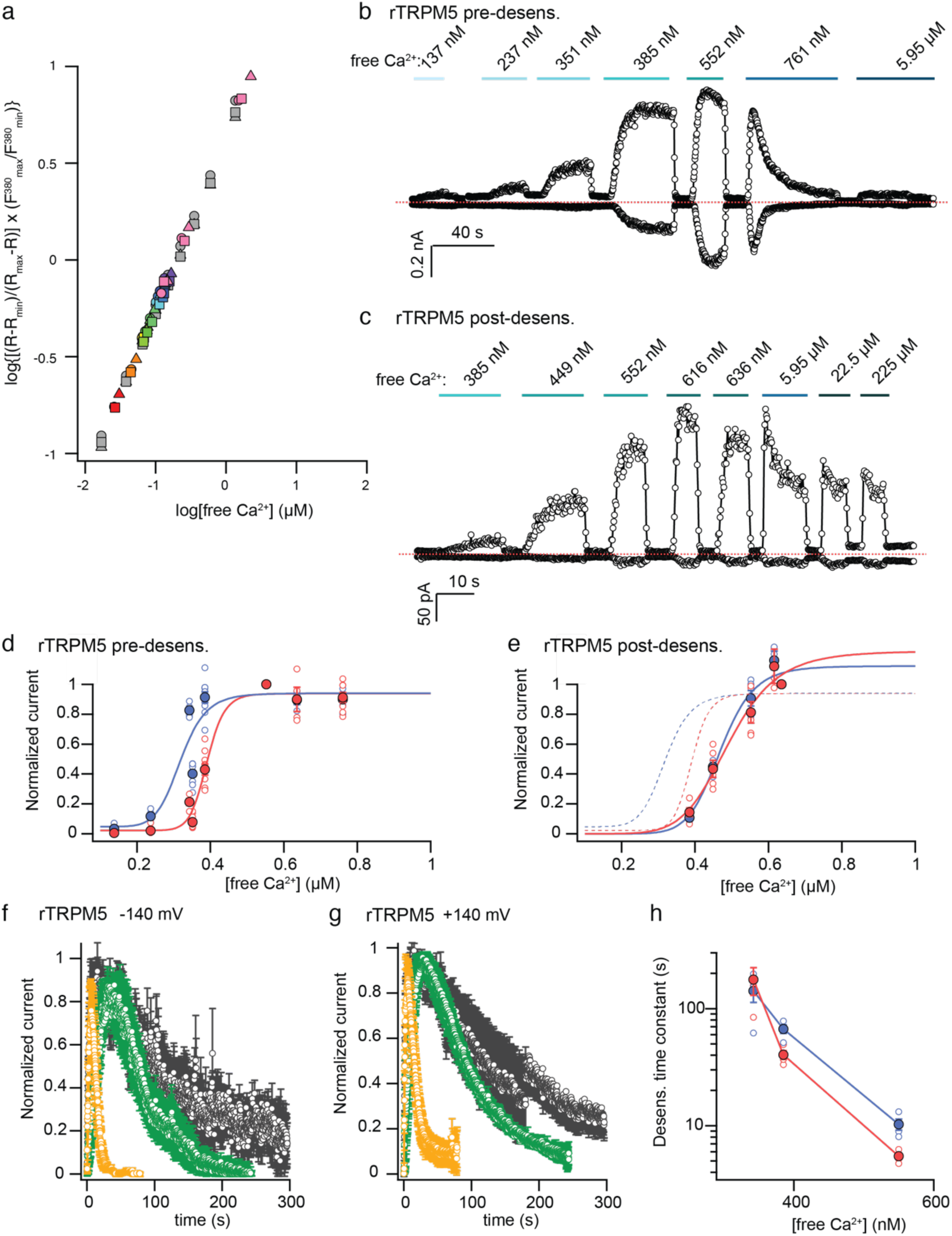
Ca^2+^-dependent de-sensitization of rTRPM5 channels. **a** Quantification using Fura-2 of the concentration of free Ca^2+^ in solutions used for patch-clamp experiments. Grey symbols are data from the standard solutions (0.017 μM, 0.038 μM, 0.065 μM, 0.1 μM, 0.225 μM, 0.351 μM, 0.602 μM, and 1.35 μM free Ca^2+^) and data for experimental solutions, diluted 1:5 as described in the Methods section, are shown in rainbow colors. The theoretical vs experimentally determined free Ca^2+^ concentrations, considering the 1:5 dilution factor, were: ● 50 nM vs 137 ± 7 nM; ● 60 nM vs 237 ± 14 nM; ● 70 nM vs 351 ± 17 nM; ● 85 nM vs 343 ± 17 nM; ● 100 nM vs 385 ±18 nM; ● 110 nM vs 449 ± 21 nM; ● 120 nM vs 552 ± 26 nM; ● 130 nM vs 616 ± 40 nM; ● 140 nM vs 636 ± 31 nM; ● 160 nM vs 761 ± 48 nM). For a sample with a theoretical concentration of 1 μM free Ca^2+^ (pink symbols), we quantified dilutions of 1:5, 1:20 and 1:50, and obtained an averaged free Ca^2+^ concentration of 5.95 ± 0.4 μM. All samples and standard solutions were prepared and measured in triplicate, with individual measurements shown as circles, triangles and squares. **b-c.** Representative rTRPM5 channel currents elicited by different concentrations of intracellular free Ca^2+^ at ± 140 mV in inside-out patches, measured (**b**) before the onset of desensitization and (**c**) after desensitization has reached steady-state in the presence of 5.95 μM free Ca^2+^. The dotted lines denote the zero-current level. **d-e** Normalized and leak-subtracted concentration-response relations for rTRPM5 channels measured **(d)** before and **(e)** after Ca^2+^- dependent desensitization, obtained as in (b) and (c) at -140 (red) or +140 mV (blue). Data is shown as mean ± SEM (before desensitization, n = 10; after desensitization, n = 5) and individual experiments are shown as open circles. Curves are fits to the Hill equation. Fit parameters were: (**d**, +140 mV) EC_50_ = 337 ± 6 nM, Hill coef. = 8.7 ± 0.4; (**d**, -140 mV) EC_50_ = 413 ± 9 nM, Hill coef. > 8; (**e**, +140 mV) EC_50_ = 474 ± 5 nM, Hill coef. = 9 ± 0.5; (**e**, -140 mV) EC_50_ = 504 ± 16.5 nM, Hill coef. = 10 ± 1.87. The dotted curves on (e) are fits to data before desensitization. **f-g.** Time courses of desensitization for rTRPM5 channels measured at (**f**) -140 mV or (**g**) +140 mV in the presence of 342 nM (grey), 385 nM (green) or 550 nM free intracellular Ca^2+^, obtained from inside-out patches. Data were leak subtracted, normalized to peak, and shown as mean ± SEM (n = 4). **h.** Desensitization time constants obtained from mono-exponential fits to experiments as in (f) and (g), at -140 mV (red) and +140 mV (blue). Data shown as mean ± SEM (filled symbols) and as individual experiments (open symbols).

**Extended data Fig. 2.**
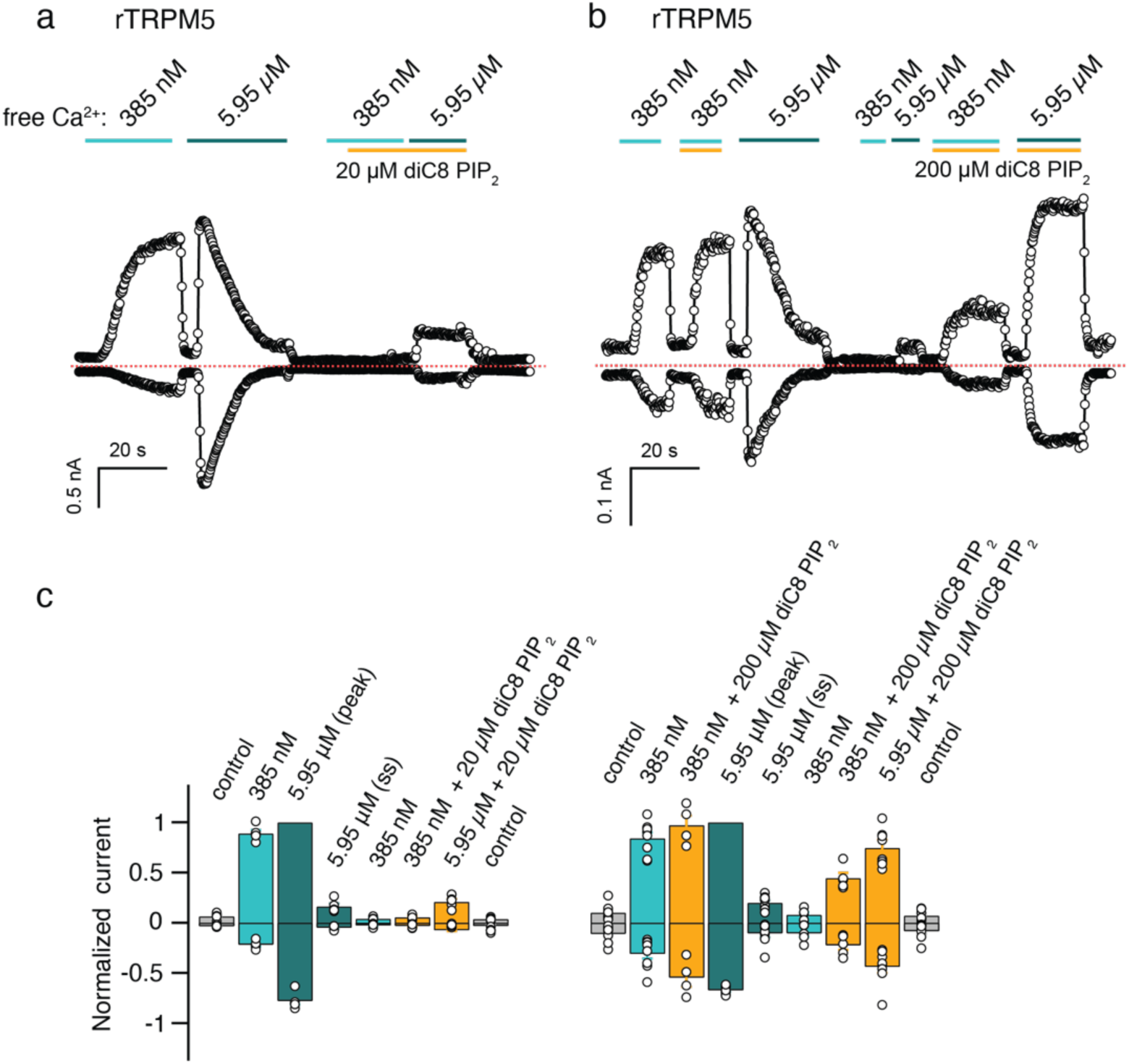
Effect of diC8 PIP_2_ on rTRPM5 channel desensitization. **a-b** Representative current time-courses at ± 140 mV in response to distinct free intracellular Ca^2+^ concentrations and (**a**) 20 or (**b**) 200 μM diC8 PIP_2_. Dotted lines denote the zero-current level. **c** Group data for experiments as in (a) and (b), depicted as mean ± SEM (n = 5 - 9) and as data from individual experiments (open circles). Data were normalized to the peak current measured in 5.95 μM free Ca^2+^ at +140 mV.

**Extended Data Fig. 3.**
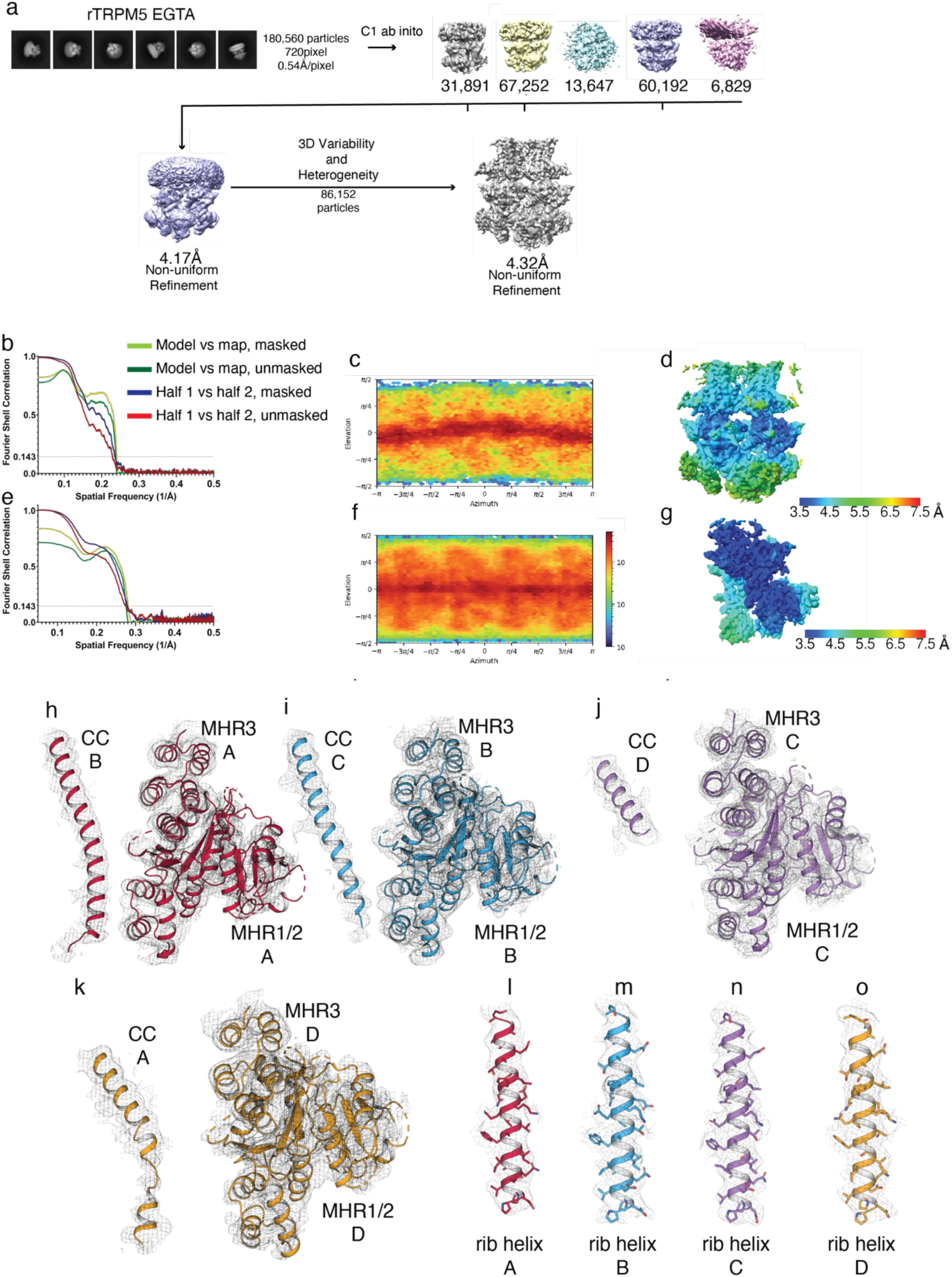
Cryo-EM data collection and processing, rTRPM5_EGTA_. **a** Data-processing workflow. **b** Fourier shell correlation calculation, **c** Euler angle distribution of particles, and **d** local resolution map of full-length protein map. **e** Fourier shell correlation calculation, **f** Euler angle distribution of particles, and **g** local resolution map of MHR generated by focused mask. **h-k** Density around CC B and MHR1/2 and MHR3 A, (interface I, **h**), CC C and MHR1/2 and MHR3 B (nterface II, **i**), CC D and MHR1/2 and MHR3 C (interface III, **j**), and CC A and MHR1/2 and MHR3 D (interface IV, **k**). All density in h-k is contoured at 0.11. **l-o** Density around rib helices in protomers A-D contoured at 0.11.

**Extended Data Fig. 4.**
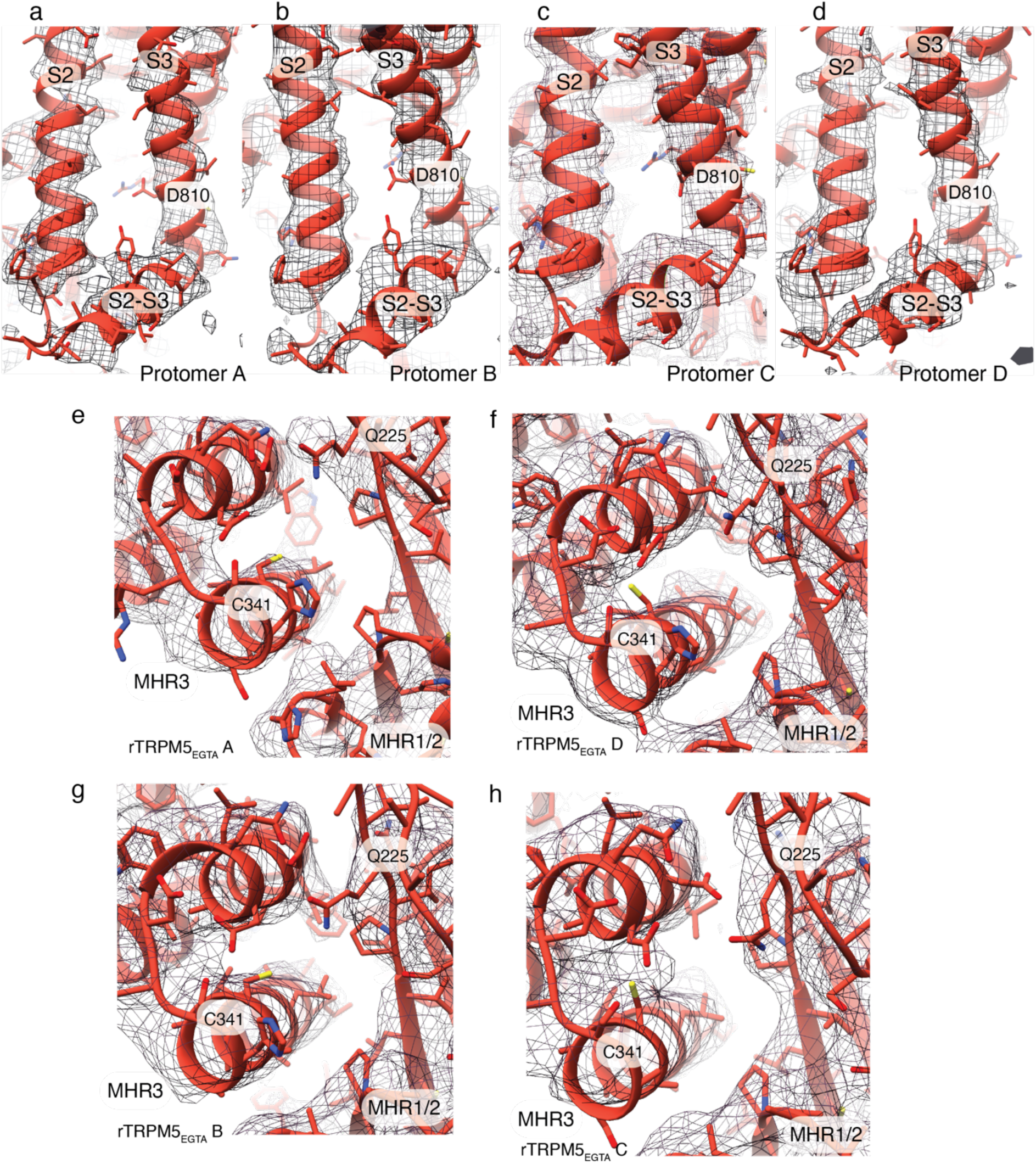
Density around the Ca^2+^ binding sites in rTRPM5_EGTA_. **a-d** The VSLD site in protomers A-D (contoured at level 0.12). **e-h** The cytoplasmic site in protomer A (**b**), D (**c**), B (**d**), and C (**e**). Density in **e-h** is contoured at level 0.12.

**Extended Data Fig. 5.**
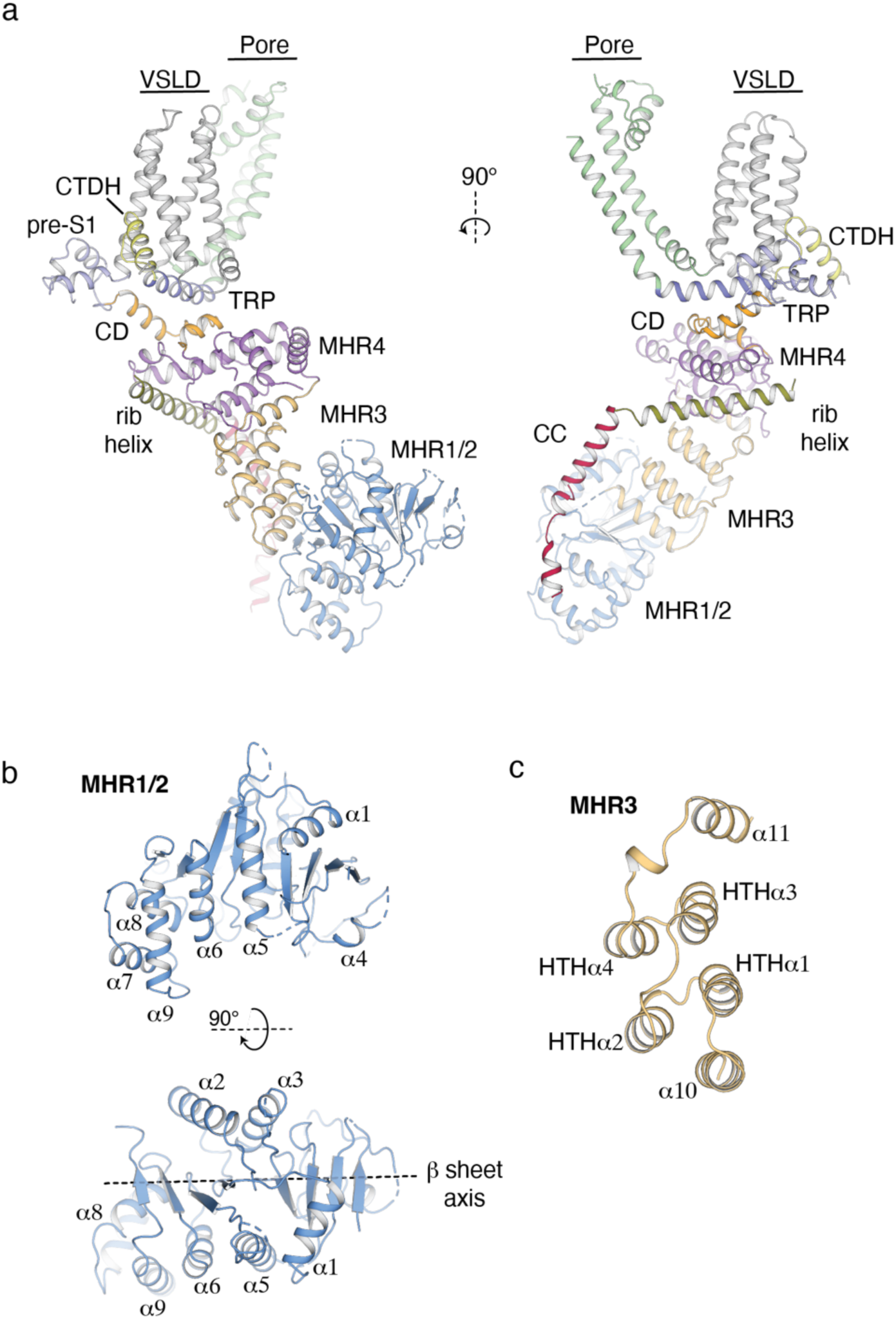
Architecture of rat TRPM5. **a** A single protomer is shown for clarity. MHR1/2 (blue), MHR3 (bright orange), MHR4 (purple), coupling domain (CD, gold), pre-S1 (lilac), Voltage sensing-like domain (VSLD, grey), pore domain (light green), TRP helix (marine blue), CTD helix (CTDH, yellow), rib helix (green), CC (red). **b** The architecture of the MHR1/2 domain. Front view (top) and top-down view (bottom). Two sets of helices sandwich a β-sheet assembly. This creates an internal β-sheet axis. **c** The MHR3 domain is made up of two helix-turn-helix (HTH) motifs flanked by individual helices above and below.

**Extended Data Fig. 6.**
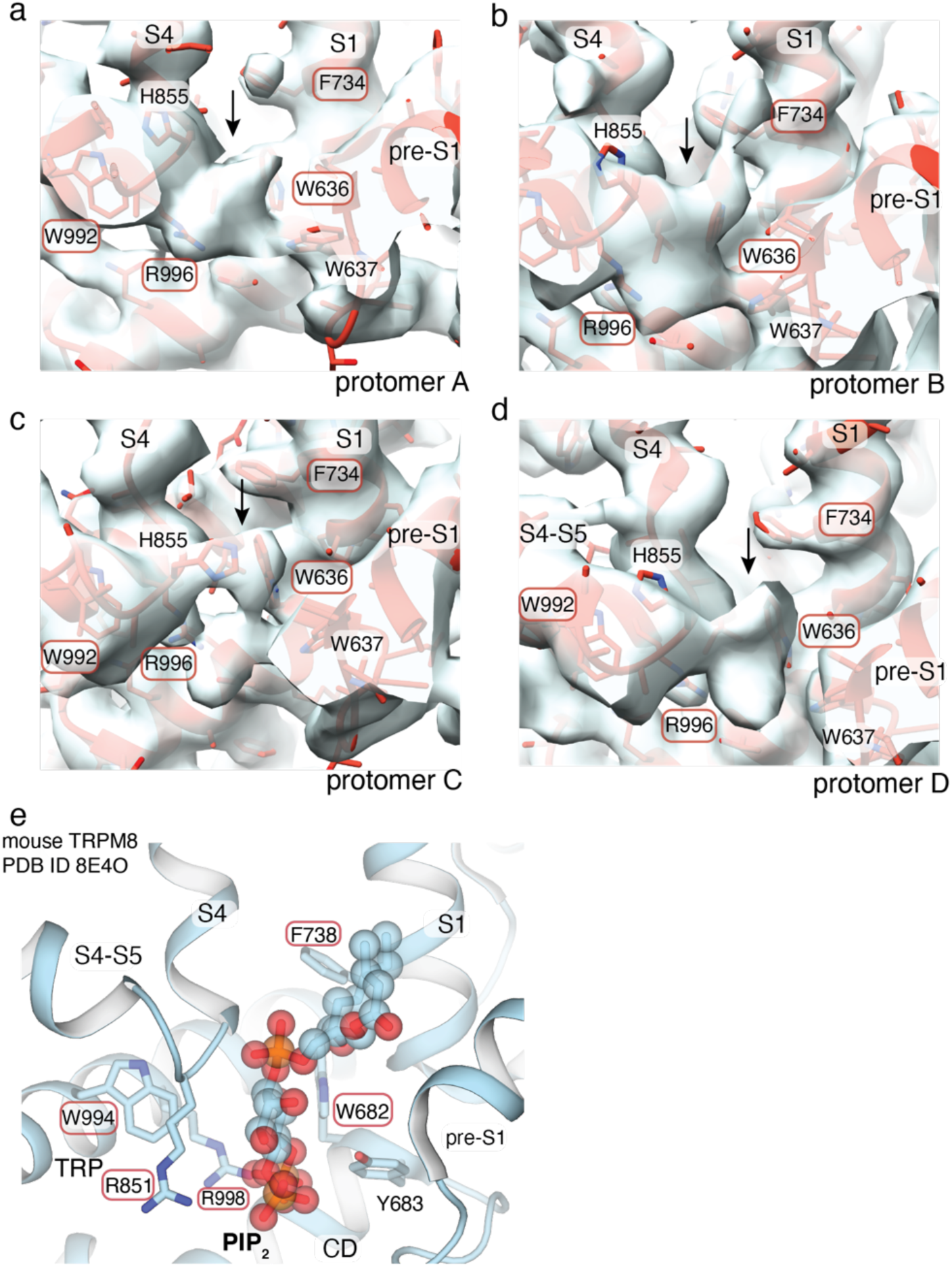
Putative PIP_2_ densities in the rTRPM5_EGTA_ structure. **a** A non-protein density observed in the cleft between pre-S1, S1, S4, S4-S5 linker, and the TRP domain in rTRPM5_EGTA_ in protomers A-D (**a**-**d**). Density is contoured at level 0.11. The site is reminiscent of the PIP_2_ binding site in mouse TRPM8. Residues conserved between rTRPM5 and mouse TRPM8 are circled in red. The density is indicated with an arrow. **e** The PIP_2_ binding site in mouse TRPM8(PDB ID 8E4O). PIP_2_ is shown in stick and sphere representation.

**Extended Data Fig. 7.**
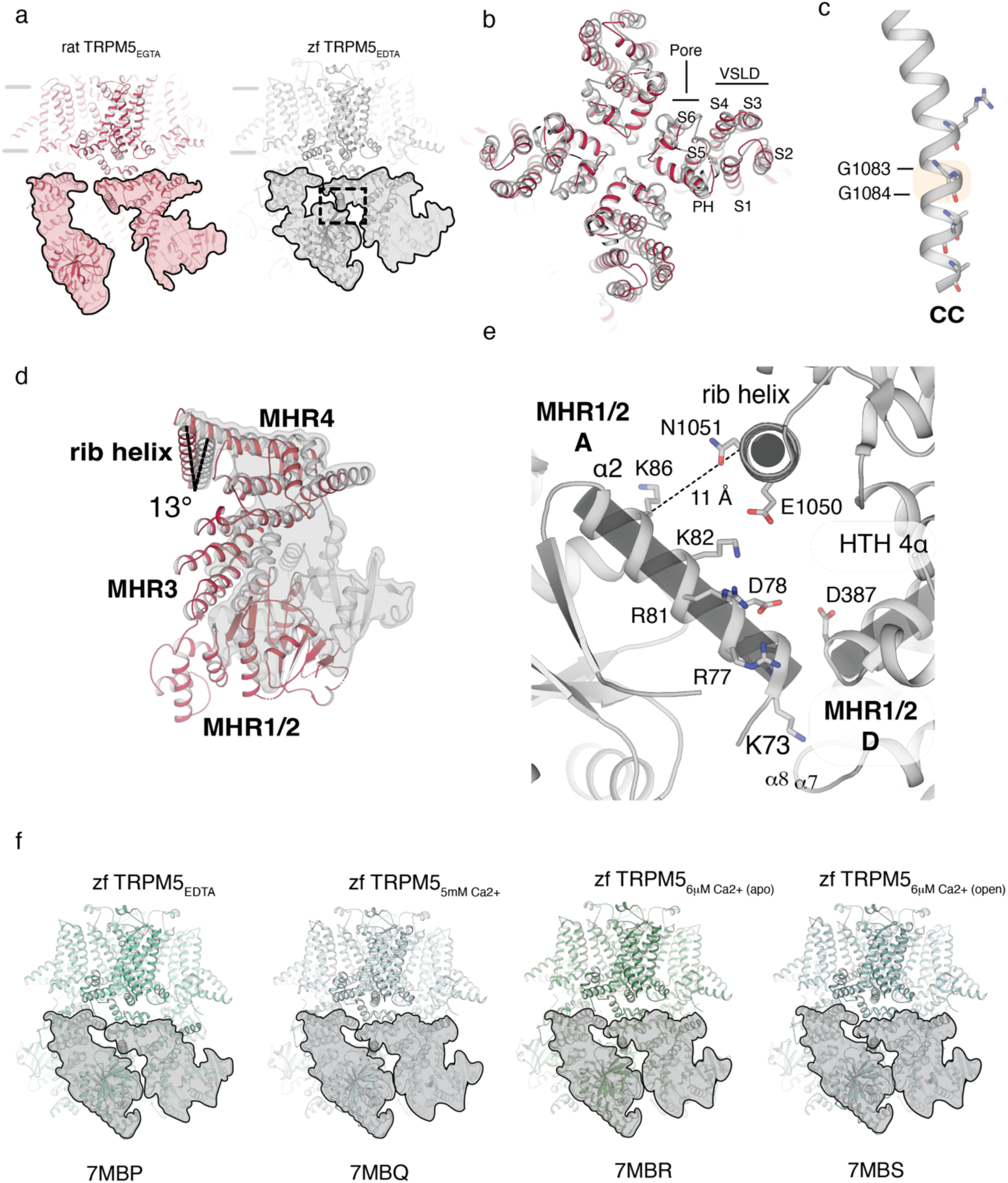
Comparison of rTRPM5_EGTA_ and zebrafish TRPM5 in the presence of EDTA (zfTRPM5_EDTA_). **a** Side-view of rTRPM5_EGTA_ (protomers A and D, as shown in red in Fig. 3a, red) and zfTRPM5_EDTA_ (grey). The cytoplasmic domains are highlighted. **b** Top view of the overlay of rTRPM5_EGTA_ (red) and zfTRPM5_EDTA_ (grey). Pore domain (S5 and S6) and Voltage Sensing-Like Domain (VSLD, S1-S4) are indicated. **c** The zfTRPM5_EDTA_ CC helix. The two glycines that make up the GG hinge (G1083 and G1084) are highlighted in an orange box. **d** Overlay of rTRPM5_EGTA_ (red) and zfTRPM5_EDTA_ (grey, highlighted). For clarity, only the cytoplasmic domains of a single protomer are shown for each channel. Protomer A is shown for rTRPM5_EGTA_. The zfTRPM5_EDTA_ is rotated by 13° around the rib helix compared to rTRPM5_EGTA_. **e** A close-up of the interprotomer interface between MHR1/2 and the rib helix, MHR3, and MHR1/2 (region indicated by dashed line box in **a**). The distance between the Cα carbons of K86 in the α2 helix of MHR1/2 and N1051 in the rib helix is 11Å. **f** Structures of zfTRPM5 captured under different Ca^2+^ conditions and in different conformations. Shown are zfTRPM5_EDTA_ (PDB ID 7MBP), zfTRPM5_5mM Ca2+_ (PDB ID 7MBQ), zfTRPM5_6μM Ca2+ (apo)_ (PDB ID 7MBR), zfTRPM5_6μM Ca2+ (open)_ (PDB ID 7MBS). The cytoplasmic domains are shaded in grey.

**Extended Data Fig. 8.**
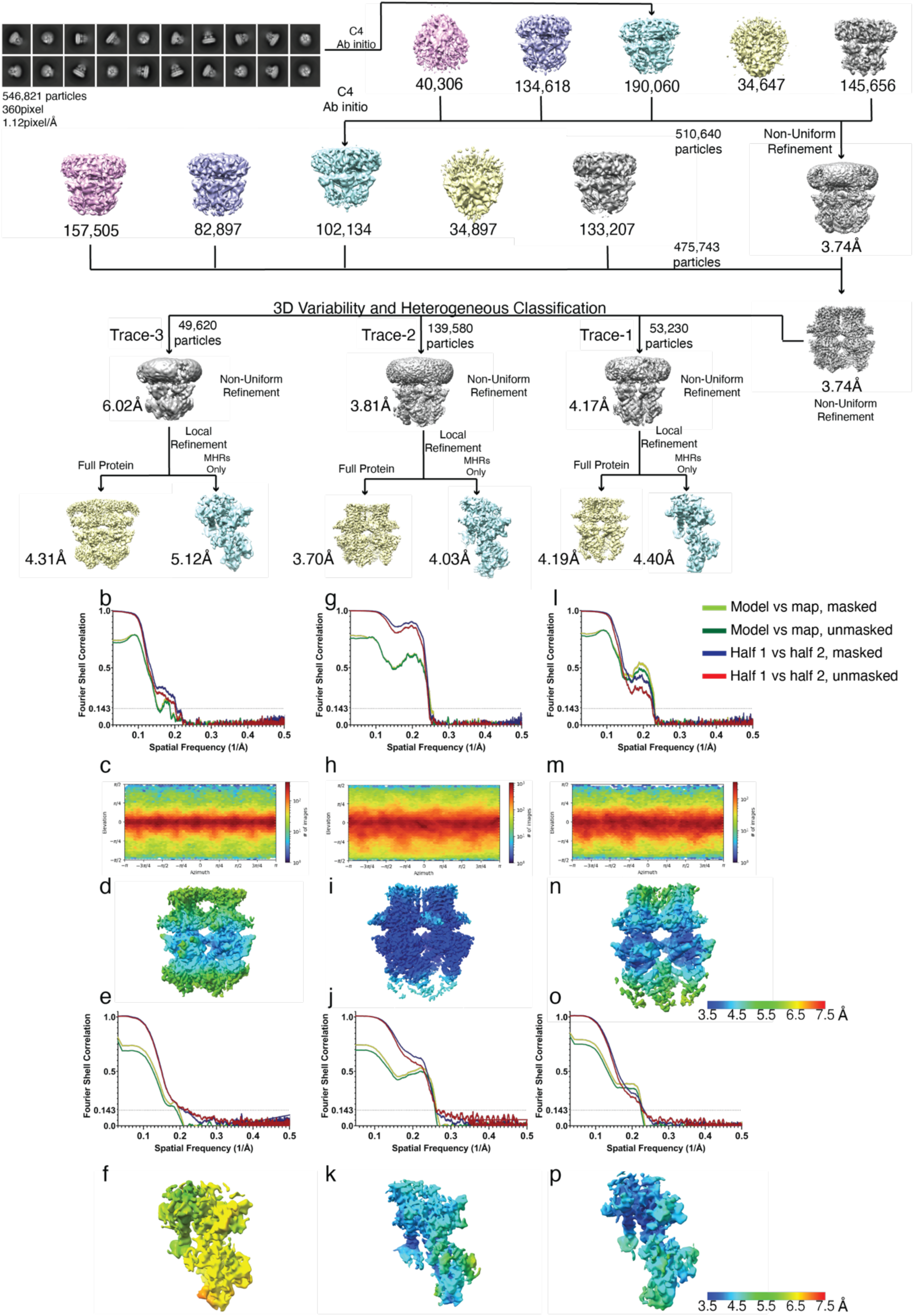
Cryo-EM processing, rTRPM5_trace_. **a** Data processing workflow of rTRPM5_trace_. **b-f** Data processing and validation for rTRPM5_trace-3_. **b** Fourier shell correlation calculation of the full-length map. **c** Euler angle distribution of particles. **d** Local resolution of full-length protein map. **e** Fourier shell correlation calculation of MHR domain map from focused masking. **f** Local resolution map of MHR domain **g-k** Data processing workflow of rTRPM5_trace-2_ **l-p** Data processing workflow of rTRPM5_trace-1._

**Extended Data Fig. 9.**
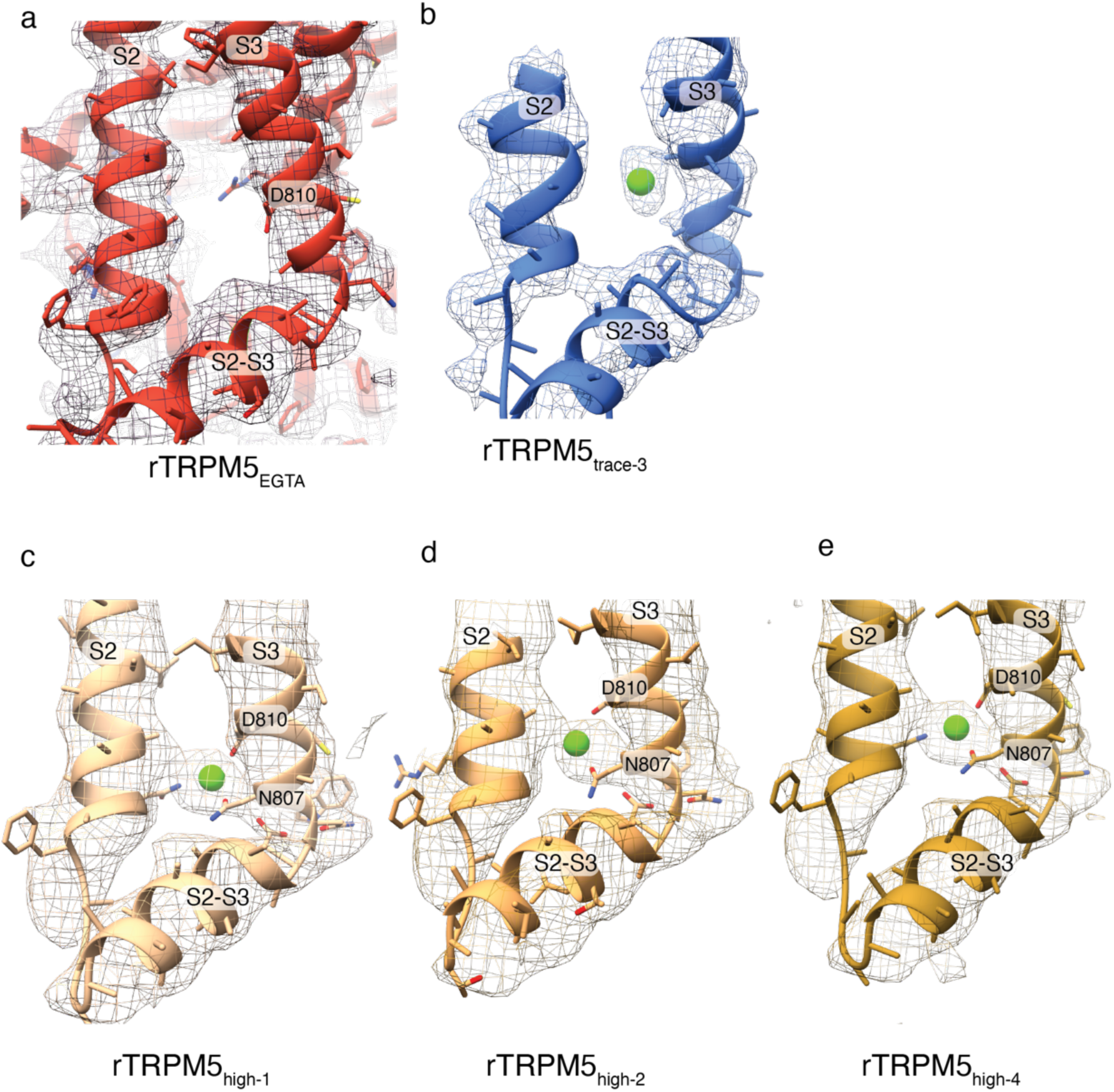
The VSLD calcium binding site in **a** rTRPM5_EGTA_ (protomer C, contoured at level 0.12), **b** rTRPM5_trace-3_ (contoured at level 0.15), **c** rTRPM5_high-1_ (contoured at level 0.15), **d** rTRPM5_high-2_ (contoured at level 0.15), and **e** rTRPM5_high-4_ (contoured at level 0.15).

**Extended Data Fig. 10.**
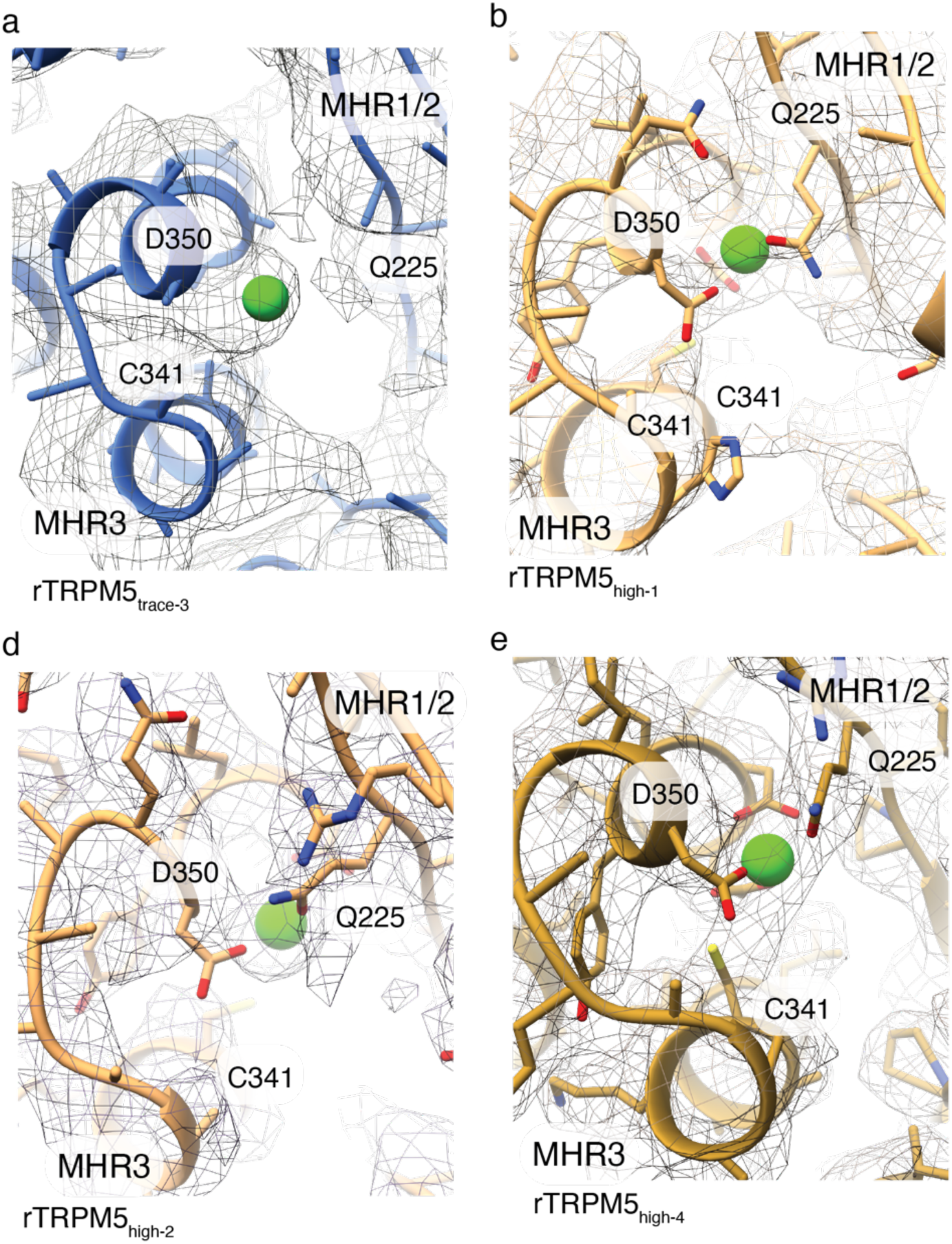
The cytosolic calcium binding site in **a** rTRPM5_trace-3_ (contoured at level 0.2), **b** rTRPM5_high-1_ (contoured at level 0.2), **c** rTRPM5_high-2_ (contoured at level 0.3), and **d** rTRPM5_high-4_ (contoured at level 0.25).

**Extended Data Fig. 11.**
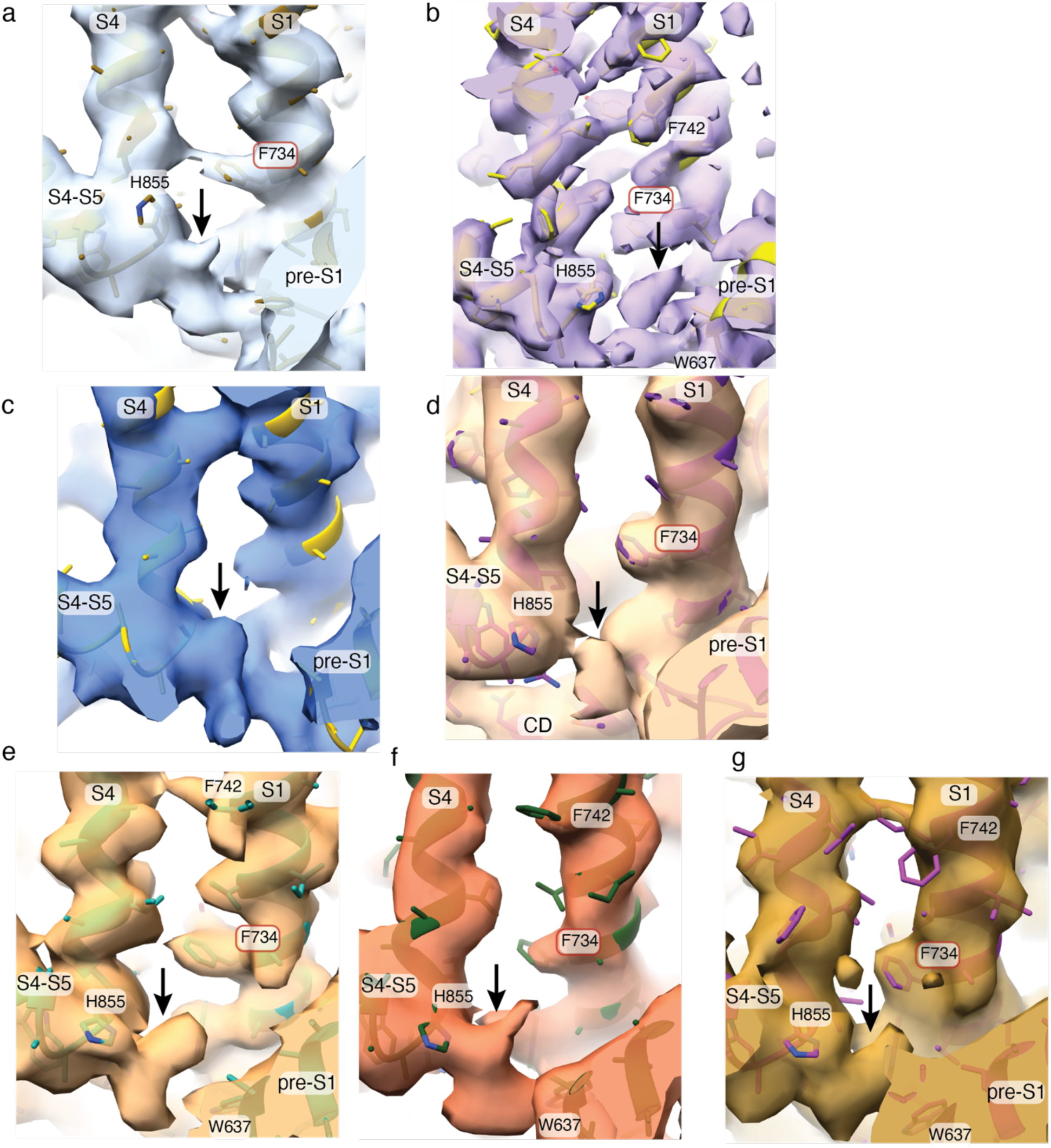
Non-protein densites observed in the putative PIP2 binding site in rTRPM5 in the presence of Ca^2+^. **a-g** Densities observed in rTRPM5_trace-1_ (**a,** contoured at level 0.16), rTRPM5_trace-2_ (**b**, contoured at level 0.2), rTRPM5_trace-3_ (**c,** contoured at level 0.15), rTRPM5_high-1_ (**d**, contoured at level 0.15), rTRPM5_high-2_ (**e**, contoured at level 0.15), rTRPM5_high-3_ (**f**, contoured at level 0.15), and rTRPM5_high-4_ (**g**, contoured at level 0.12).

**Extended Data Fig. 12.**
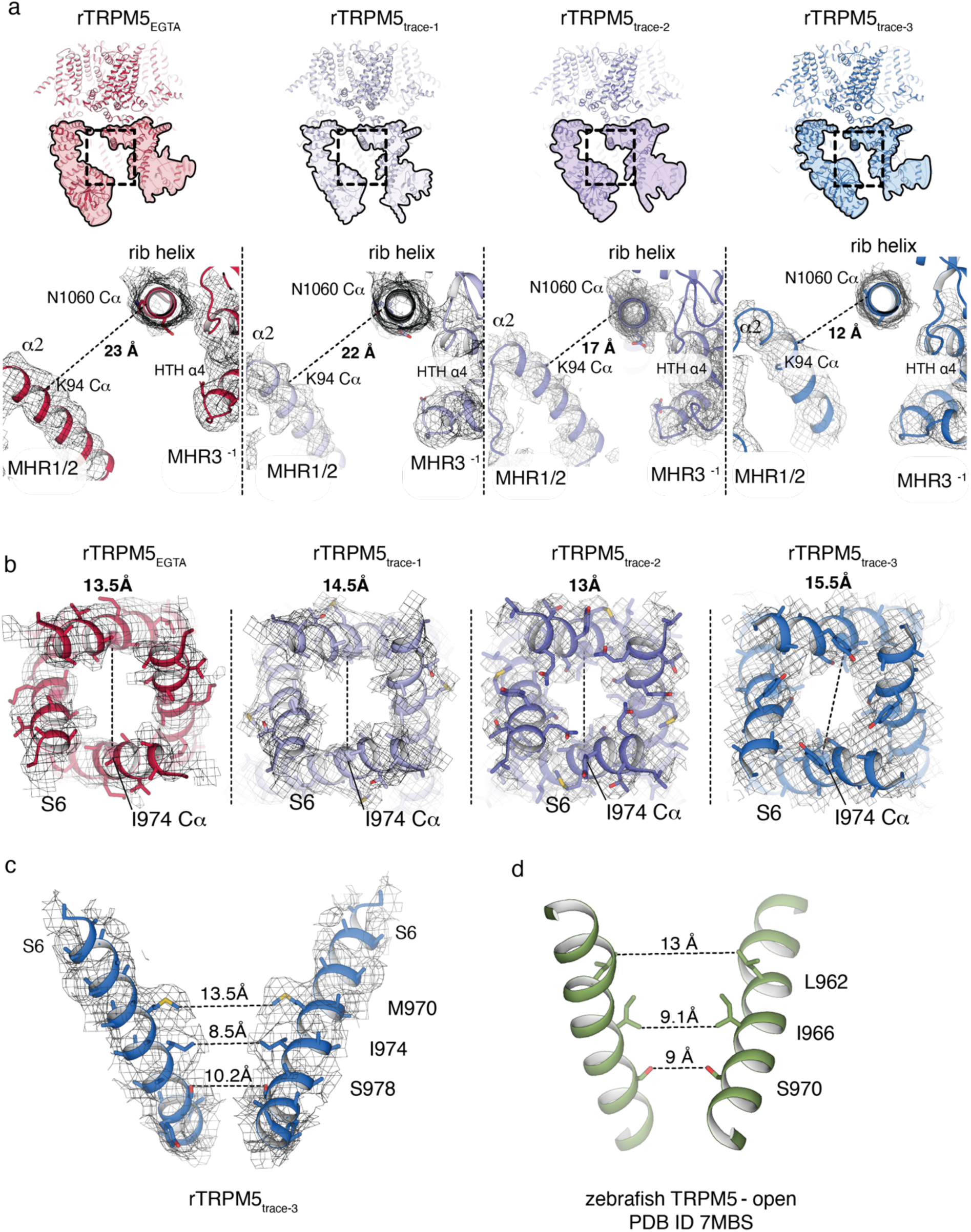
The interprotomer interfaces and pore conformations in rTRPM5_trace_ structures. **a** The boxed region in the top panel indicates the interface between MHR1/2 and the rib helix and MHR3 of neighboring protomers. The bottom panel shows a close-up of the boxed region in rTRPM5_EGTA_ (protomers A and D, red), rTRPM5_trace-1_ (lilac), rTRPM5_trace-2_ (purple), and rTRPM5_trace-3_ (blue). Density is shown for the interface in all structures and contoured at levels: 0.12 (rTRPM5_EGTA_), 0.13 (rTRPM5_trace-1_), 0.14 (rTRPM5_trace-2_), and 0.15 (rTRPM5_trace-3_). Distance is measured between Cα carbons of K94, located in the MHR1/2 helix α2, and N1060 located in the rib helix. This distance decreases from 22 Å in rTRPM5_trace-1_ to 10 Å in rTRPM5_trace-3_. **b** Comparison of pore conformations in rTRPM5_EGTA_ (red), rTRPM5_trace-1_ (lilac), rTRPM5_trace-2_ (purple), and rTRPM5_trace-3_ (blue). The bottom-up view is shown. Density is shown for the S6 helix in all structures. Distance is measured between Cα carbons of the gate residue I974 (shown in spheres) of opposing protomers. S6 side chains are shown in stick representation where the y have been built. The density is contoured at levels: 0.1 (rTRPM5_EGTA_), 0.12 (rTRPM5_trace-1_), 0.15 (rTRPM5_trace-2_), and 0.1 (rTRPM5_trace-3_) **c** Side-view of the rTRPM5_trace-3_ pore and the corresponding density contoured at level 0.1, with only two pore-lining S6 helices shown for clarity. Side chains are shown for residues M970, I974, S978, and Y979. **d** The pore of the zebrafish TRPM5 channel in an open conformation (PDB ID 7MBS) for comparison. The residues L962, I966, and S970 are shown in stick representation.

**Extended Data Fig. 13.**
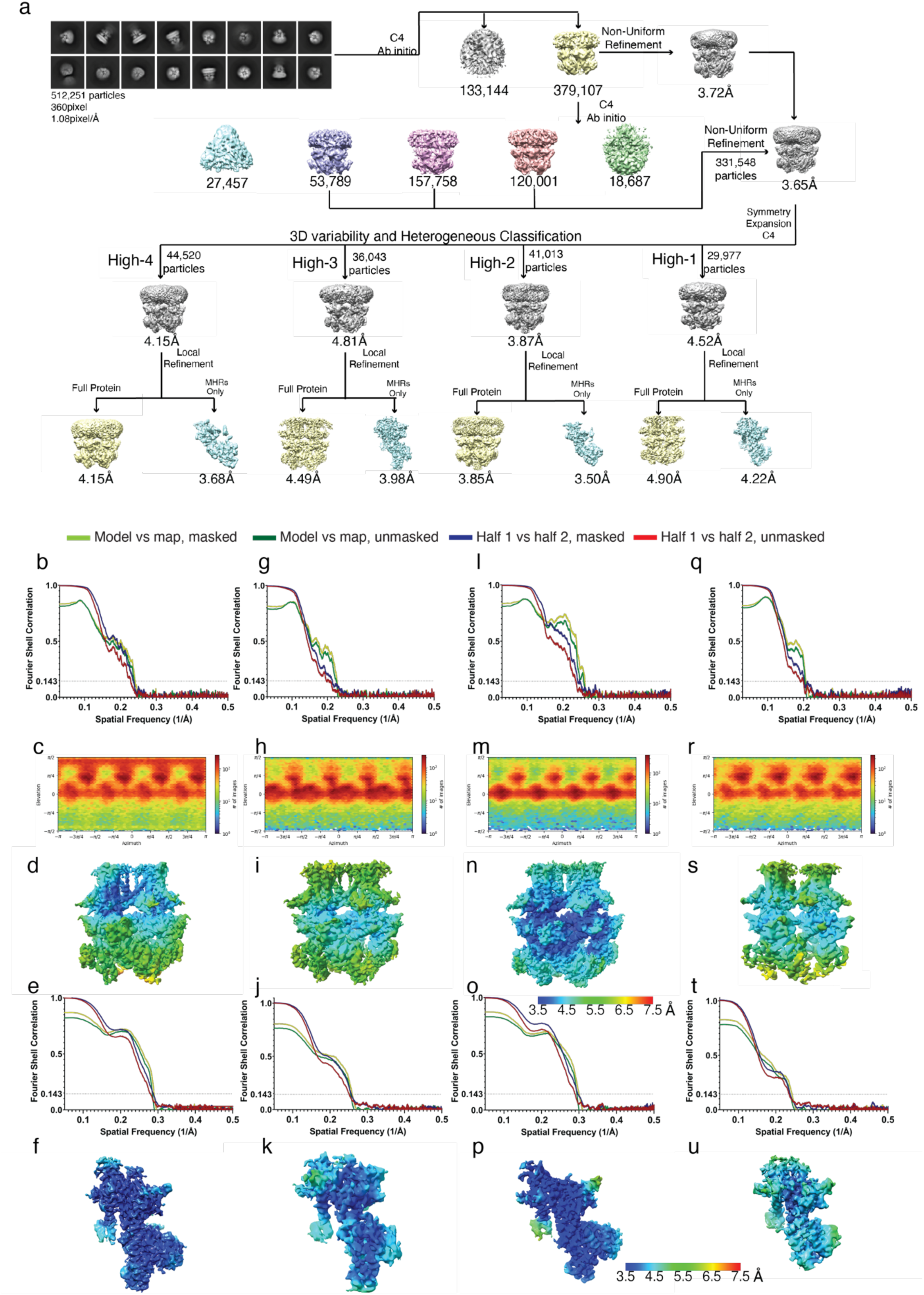
Cryo-em workflow for therTRPM5_high_ dataset. **a** data processing workflow of rTRPM5_high_. **b-f** Processing and validation for rTRPM5_high-4._ **b** Fourier shell correlation calculation of the full-length map. **c** Euler angle distribution of particles. **d** Local resolution of full-length protein map. **e** Fourier shell correlation calculation of MHR domain map from focused masking. **f** Local resolution map of MHR domain from focused masking. **g-k** Processing and validation for rTRPM5_high-3_. **l-p** Processing and validation for rTRPM5_high-2._ **q-u** Processing and validation for rTRPM5_high-1._

**Extended Data Fig. 14.**
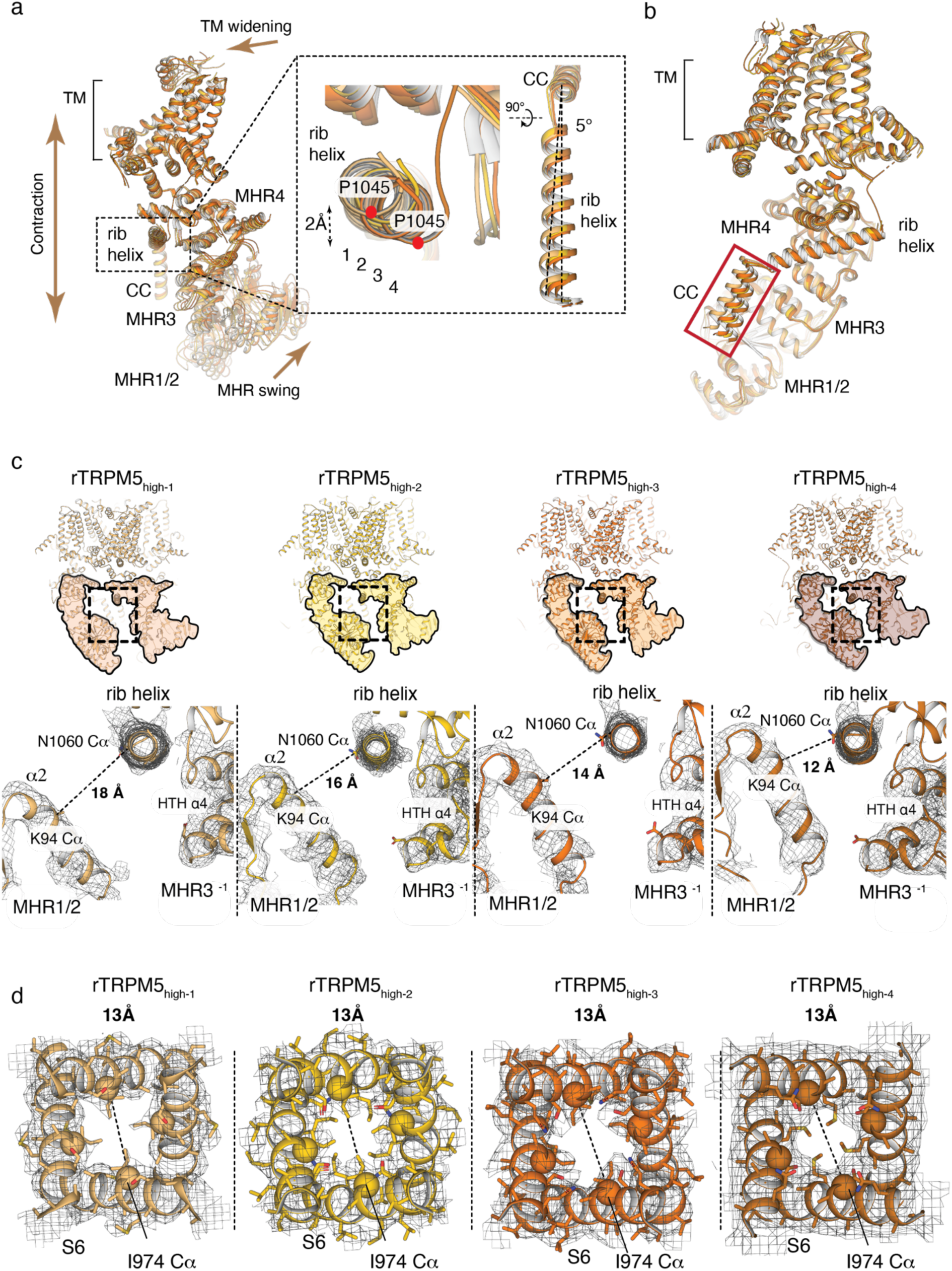
The interprotomer interfaces and pore conformations in rTRPM5_trace_ structures. **a** Alignment of rTRPM5_high-1_ (bright orange), rTRPM5_high-2_ (gold), rTRPM5_high-3_ (deep orange), and rTRPM5_high-4_ (maroon). For clarity, only a single protomer is shown for each structure. As previously observed in rTRPM5_trace_ structures, a rotation occurs around the rib helix (indicated by box and enlarged inset). The rib helix of rTRPM5_high-4_ is rotated by 5° and displaced laterally by ∼2 Å compared to rTRPM5_high-1_. The red dot signifies the position of P1045 in the rib helix. **b** An overlay of the individual rTRPM5_high-1-4_ protomers shown in **a**. The superposition results in good agreement between the protomers in all regions except in the CC (red box). **c** Comparison of inter-protomer interfaces in rTRPM5_high-1-4 ._ Boxed region in the top panel indicates the interface between MHR1/2 and the rib helix and MHR3 of neighboring protomers. The bottom panel shows a close-up of the boxed region in rTRPM5_high-1_, rTRPM5_high-2_, rTRPM5_high-3_, and rTRPM5_high-4._ Density is shown for the interface in all structures. It is contoured at level 0.1 (rTRPM5_high-1_), 0.12 (rTRPM5_high-2_), 0.1 (rTRPM5_high-3),_ and 0.15 (rTRPM5_high-_4). Distance is measured between Cα carbons of K94, located in the MHR1/2 helix α2, and N1060 located in the rib helix. **d** Comparison of pore conformations in rTRPM5_high-1_, rTRPM5_high-2_, rTRPM5_high-3_, and rTRPM5_high-4._ The bottom-up view is shown. Density is shown for the S6 helix in all structures, contoured at level 0.12 (rTRPM5_high-1_), 0.15 (rTRPM5_high-2_), 0.1 (rTRPM5_high-3),_ and 0.1 (rTRPM5_high-_4). Distance is measured between Cα carbons of the gate residue I974 (shown in spheres) of opposing protomers. Side chains of S6 residues are shown in stick representation where they have been built.

**Extended Data Fig. 15.**
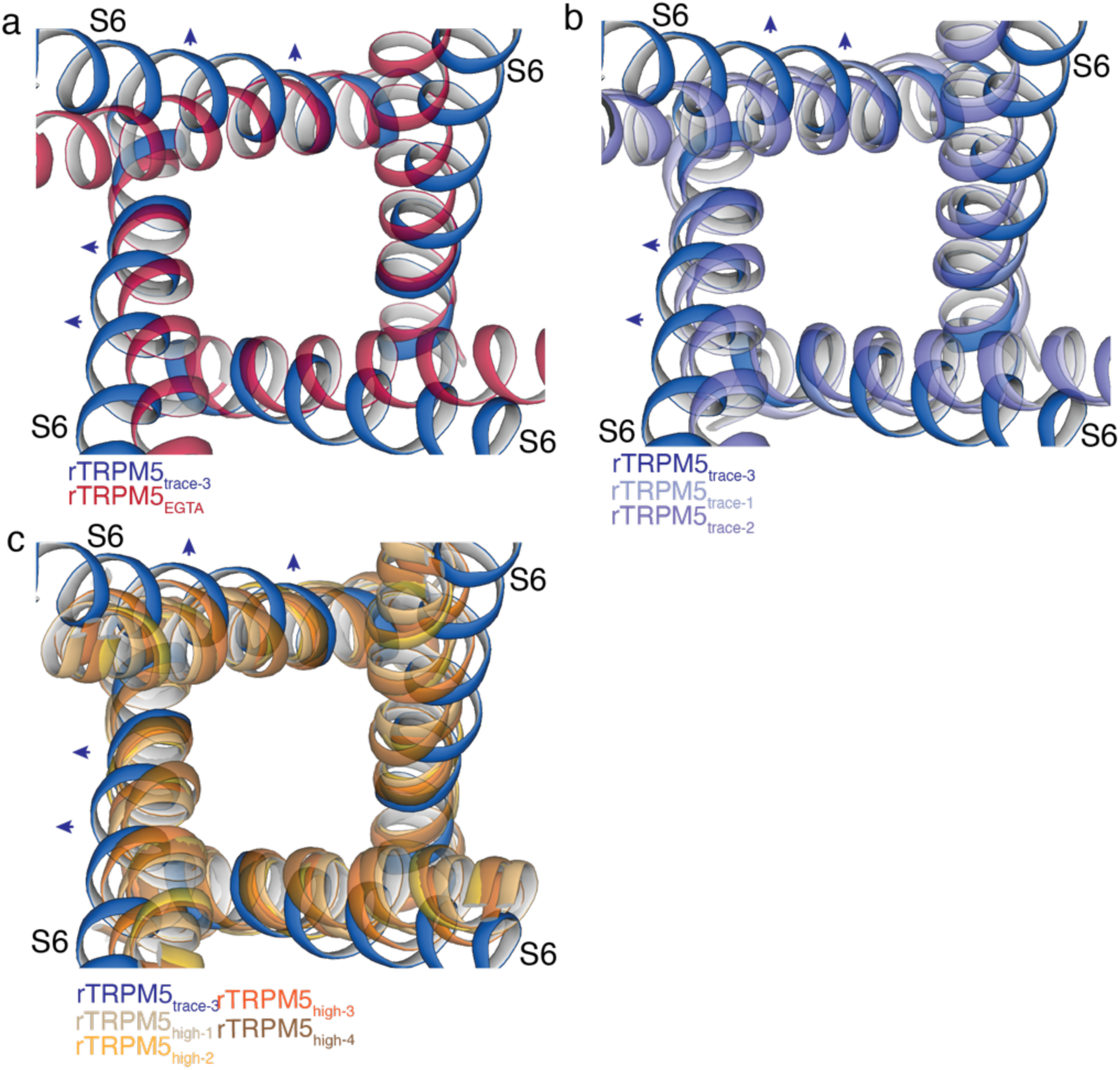
Pore dimensions in rTRPM5 structures. **a-c** Top-down view of the overlay of the rTRPM5_trace-3_ pore (blue, tentatively open) with (**a**) rTRPM5_EGTA_ (red), (**b**) rTRPM5_trace-1_ (lilac) and rTRPM5_trace-2_ (purple), (**c**) rTRPM5_high-1_ (light orange), rTRPM5_high-2_ (gold), rTRPM5_high-3_ (deep orange), and rTRPM5_high-3_ (maroon). The rTRPM5_trace-3_ pore is wider compared to the others.

**Extended Data Fig. 16.**
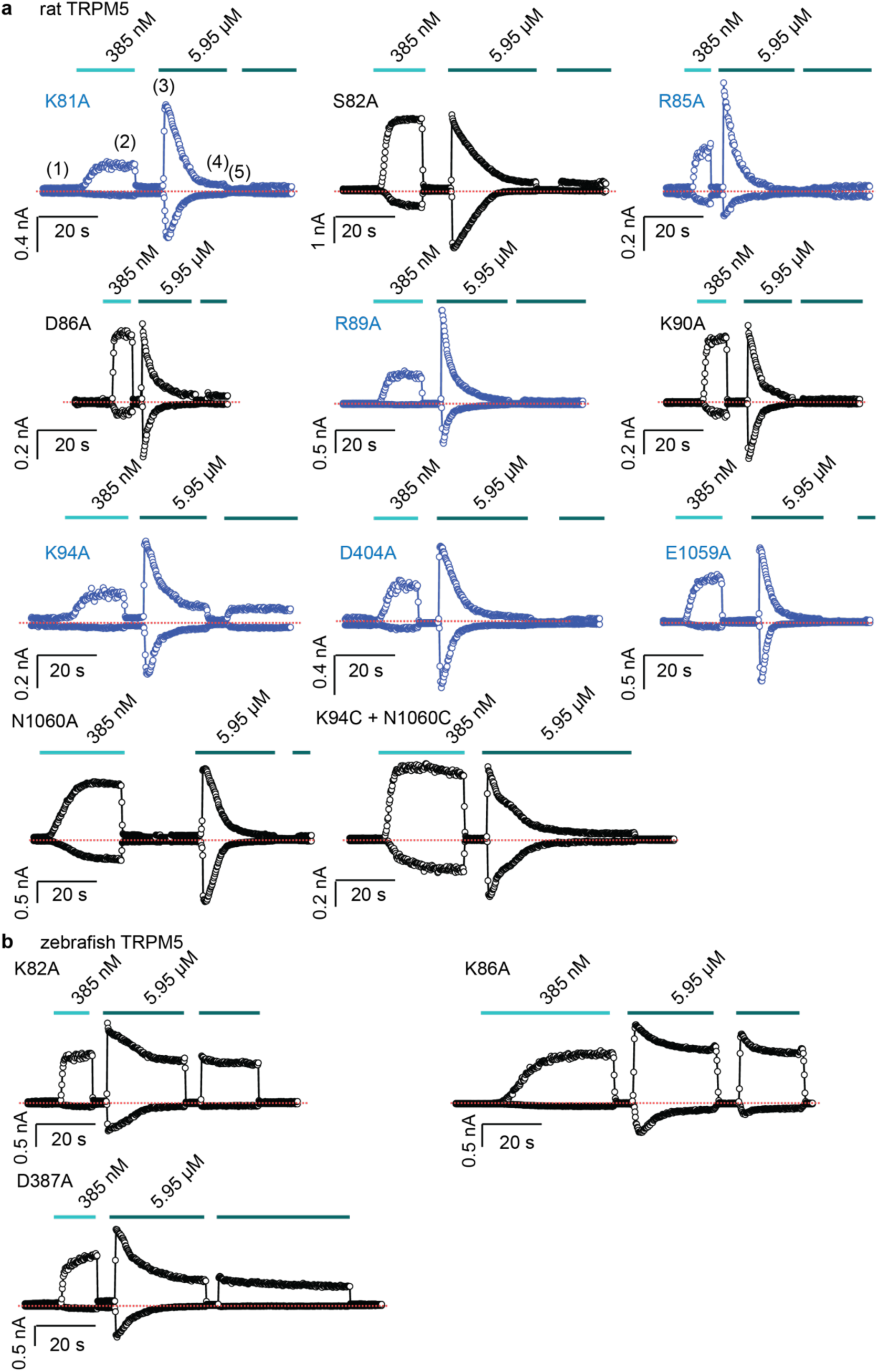
Current time-courses from mutant rTRPM5 channels. **a** Representative current time-courses at ± 140 mV obtained from inside-out patches from cells expressing rat TRPM5 channel mutants. Dotted lines denote the zero-current level. Traces in black are from mutants with WT-like behavior, and traces in blue are from mutants with a reduced response to 385 nM free Ca^2+^. Group data shown in Fig. 7e. **b** Representative current time-courses at ± 140 mV obtained from inside-out patches from cells expressing mutant zebrafish TRPM5 channels. Group data shown in Fig. 7f.

**Extended Data Fig. 17.**
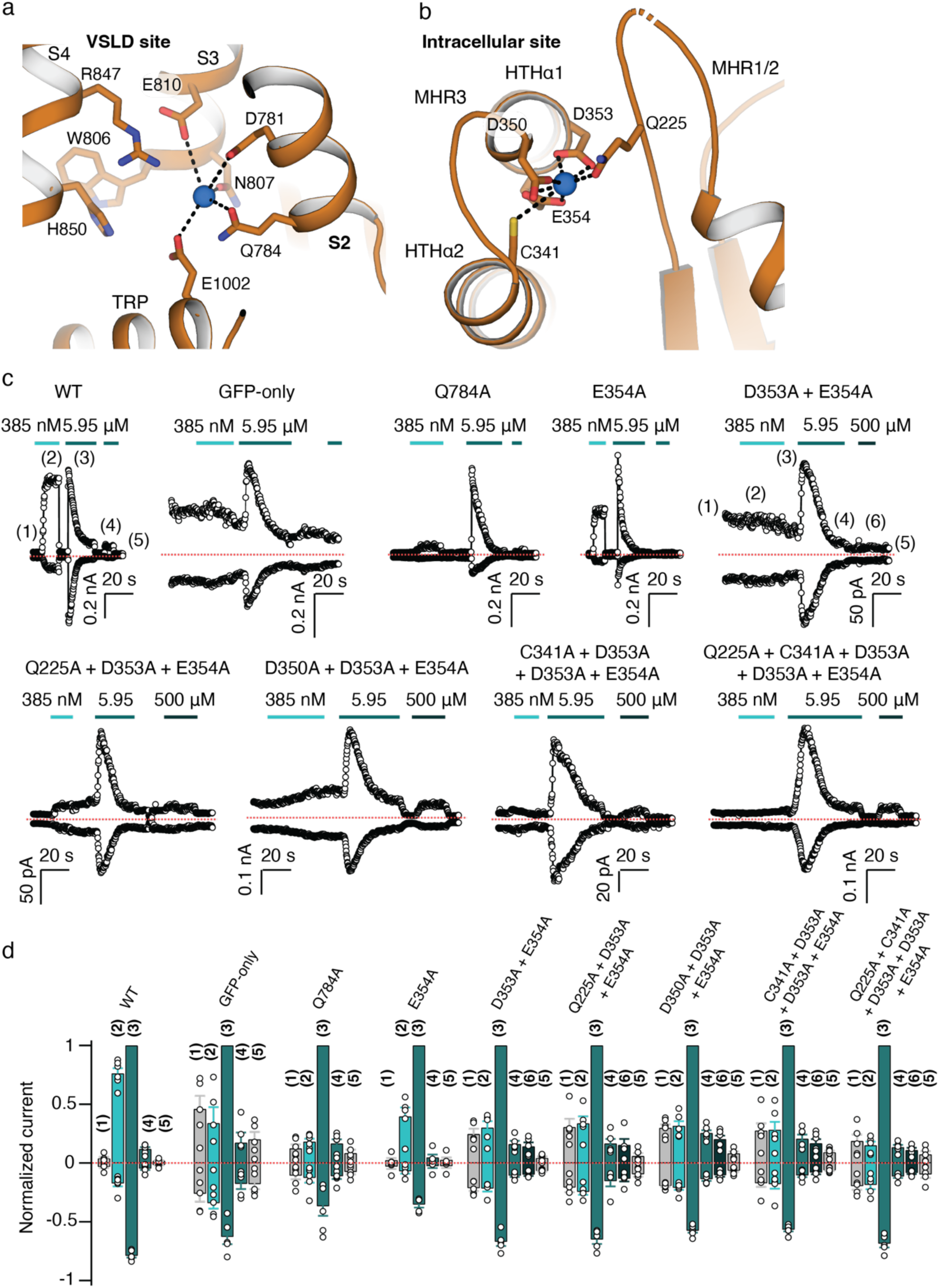
Calcium-binding site mutants in rTRPM5. **a** The VSLD Ca^2+^ binding site shown in rTRPM5_high-4_. **b** The intracellular Ca^2+^ binding site shown in rTRPM5_high-4_. **c** Representative current time-courses at ± 140 mV obtained from inside-out patches from cells transfected with GFP only, or GFP together with WT or mutant rTRPM5 channel cDNA. Dotted lines denote the zero-current level. The numbers in parenthesis on the WT-time-course denote the following: (1 and 5) zero free Ca^2+^ control solution at the beginning and end of experiment; (2) steady-state current at 385 nM free Ca^2+^ before desensitization; (3) peak current at 5.95 μM free Ca^2+^; (4) steady-state current at 5.95 μM free Ca^2+^ after desensitization; (6) The steady state currents at 500 μM free Ca^2+^ were measured after desensitization for some of the mutant rTRPM5 channels. **d** Group data for experiments in (c), shown as mean ± SEM (n = 4 – 6) or as individual experiments (open circles). Data were normalized to the peak current at +140 mV and 5.95 μM free Ca^2+^. Numbers in parenthesis denote measurements under the conditions described above.

**Table 1:** Cryo-EM data collection, refinement and validation statistics.

|  | rTRPM5<sup>EGTA</sup><br>(EMDB-40574)<br>(PDB 8SL6) | rTRPM5<sup>trace-</sup><sub>1</sub><br>(EMDB-40575)<br>(PDB 8SL8) | rTRPM5<sup>trace-</sup><sub>2</sub><br>(EMDB-40576)<br>(PDB 8SLA) | rTRPM5<sup>trace-</sup><sub>3</sub><br>(EMDB-40577)<br>(PDB 8SLE) | rTRPM5<sup>high-</sup><sub>1</sub><br>(EMDB-40578)<br>(PDB 8SLI) | rTRPM5<sup>high-</sup><sub>2</sub><br>(EMDB-40579)<br>(PDB 8SLP) | rTRPM5<sup>high-</sup><sub>3</sub><br>(EMDB-40580)<br>(PDB 8SLQ) | rTRPM5<sup>high-</sup><sub>4</sub><br>(EMDB-40581)<br>(PDB 8SLW) |
| --- | --- | --- | --- | --- | --- | --- | --- | --- |
| **Data collection and processing** |  |  |  |  |  |  |  |  |
| Magnification | 81,000 | 81,000 | 81,000 | 81,000 | 81,000 | 81,000 | 81,000 | 81,000 |
| Voltage (kV) | 300 | 300 | 300 | 300 | 300 | 300 | 300 | 300 |
| Electron exposure (e<sup>-</sup>/Å<sup>2</sup>) | 40 | 45 | 45 | 45 | 43.3 | 43.3 | 43.3 | 43.3 |
| Defocus range (μm) | -1.0 to -2.25 | -1.0 to -2.5 | -1.0 to -2.5 | -1.0 to -2.5 | -1.0 to -2.25 | -1.0 to -2.25 | -1.0 to -2.25 | -1.0 to -2.25 |
| Pixel size (Å) | 1.08 | 1.08 | 1.08 | 1.08 | 1.12 | 1.12 | 1.12 | 1.12 |
| Symmetry imposed | C1 | C4 | C4 | C4 | C4 | C4 | C4 | C4 |
| Initial particle images (no.) | 1,462,359 | 2,601,822 | 2,601,822 | 2,601,822 | 5,766,941 | 5,766,941 | 5,766,941 | 5,766,941 |
| Final particle images (no.) | 86,152 | 52,230 | 139,580 | 49,620 | 29,977 | 41,013 | 36,043 | 44,520 |
| Map resolution (Å) | 4.4 | 4.2 | 3.7 | 4.3 | 4.9 | 3.8 | 4.5 | 4.2 |
| **Refinement** |  |  |  |  |  |  |  |  |
| Initial model used | 7MBP | this study | this study | this study | this study | this study | this study | this study |
| Map sharpening *B* factor (Å<sup>2</sup>) | -133.5 | -123.6 | -116.0 | -171.4 | -216.3 | -115.5 | -184.6 | -125.3 |
| Model composition |  |  |  |  |  |  |  |  |
| Non-hydrogen atoms | 23,955 | 19,688 | 24,500 | 17,388 | 20,988 | 24,348 | 22,916 | 25,956 |
| Protein residues | 3,634 | 3,152 | 3,608 | 3,468 | 3,576 | 3,640 | 3,624 | 3,684 |
| R.m.s. deviations |  |  |  |  |  |  |  |  |
| Bond lengths (Å) | 0.005 | 0.004 | 0.004 | 0.018 | 0.004 | 0.003 | 0.003 | 0.005 |
| Bond angles (°) | 0.690 | 0.666 | 0.650 | 0.887 | 0.568 | 0.554 | 0.642 | 0.732 |
| **Validation** |  |  |  |  |  |  |  |  |
| MolProbity score | 1.9 | 1.77 | 1.87 | 1.90 | 2.00 | 1.71 | 1.80 | 1.92 |
| Clashscore | 11 | 10 | 10 | 9 | 11 | 7 | 11 | 14 |
| Poor rotamers (%) | 0 | 0.0 | 0.0 | 0.0 | 0.0 | 0.0 | 0.0 | 0.0 |
| Ramachandran plot |  |  |  |  |  |  |  |  |
| Favored (%) | 96 | 96.3 | 95.4 | 93.6 | 93.5 | 95.7 | 96.3 | 96.0 |
| Allowed (%) | 4 | 3.7 | 4.6 | 6.4 | 6.5 | 4.3 | 3.7 | 4.0 |
| Disallowed (%) | 0.0 | 0.0 | 0.0 | 0.0 | 0.0 | 0.0 | 0.0 | 0.0 |

## Notes

### Competing Interest Statement

The authors have declared no competing interest.

